# Trash-basket epiphytes as secondary foundation species: a review of their distribution and effects on biodiversity and ecosystem functions

**DOI:** 10.1101/2021.06.22.449473

**Authors:** Gabriel Ortega-Solis, Iván A. Díaz, Daniela Mellado-Mansilla, Camila Tejo, Francisco Tello, Dylan Craven, Holger Kreft, Juan J. Armesto

## Abstract

**Background:** Secondary foundation species (FS) are organisms that inhabit ecosystems structurally defined by a primary foundation species, providing additional structure to habitats and communities. Trash-basket epiphytes (TBE) are secondary FS that enhance arboreal soil accumulation, providing shelter to animals, and rooting sites for plants. While their importance may vary across biomes, TBE have been overlooked as drivers of biodiversity and ecosystem functions. Here, we discuss the prevalence of TBE across biomes, their effects on biodiversity and ecosystem functions, and future research directions.

**Methods:** We performed a systematic literature review of articles, books and theses and collated and synthesised information about the taxonomic distribution of TBE, their effects on ecosystem functions, and reports of plant-animal and plant-plant interactions. Then, we analysed the global distribution of TBE using a generalized linear model and summarised two studies to assess their effects on soil invertebrates.

**Results:** We identified 120 publications describing 209 species of TBE. Most TBE belong to Araceae (43%), Polypodiaceae (23%), and Orchidaceae (14%) and occur in all tropical and southern temperate forests. TBE richness peaks in the South-American Pacific mangroves, Eastern Cordillera Real, and the Napo moist forests. TBE effects on ecosystem functions include arboreal soil accumulation, water retention and temperature regulation in the canopy, and nutrient leaching through stem-flow. TBE provide shelter to species in more than 97 animal families, including from invertebrates to mammals, while 72 vascular plants have been reported to root in arboreal soil of TBE.

**Conclusions:** TBE are a compelling group of model organisms that can be used to study ecological processes such as facilitation cascades, niche construction, extended phenotypes, or the effects of secondary FS on biodiversity and ecosystem functioning. TBE should also be included in forest management plans to enhance the availability of microhabitats in the canopy supporting its associated flora and fauna.

## INTRODUCTION

Foundation species (FS; Dayton 1972) are organisms that *dominate an assemblage numerically and in overall size (usually mass), determine the diversity of associated taxa through non-trophic interactions, and modulate fluxes of nutrients and energy at multiple control points in the ecosystem it defines (Ellison 2019).* This concept is closely related to keystone species (Holling 1992) and ecosystem engineers (Jones *et al*. 1994). Primary FS like trees (Ellison *et al*. 2010), corals (Graham 2014), or seagrass (Thomson *et al*. 2015) have been widely studied. However, secondary FS are also found in ecosystems shaped by a primary FS. This group of secondary FS include seaweeds, barnacles, and epiphytes (Angelini and Silliman 2014; Thomsen *et al*. 2018). The interaction between primary and secondary FS enhances species richness and abundance in an ecosystem primarily via niche construction (Thomsen *et al*. 2018).

Trash-basket epiphytes (TBE; Benzing 1990) can be classified as secondary FS in forest ecosystems. TBE are litter-trapping plants (*sensu* Zona and Christenhusz 2015) that dwell on trees and possess specialized morphological adaptations to capture significant amounts of litter and debris that is further decomposed – together with the old roots and senescent leaves of the TBE – and incorporated into arboreal soil. TBE share the litter trapping role in forest canopies with tank bromeliads, but they differ because tank bromeliads are highly specialized in retaining water as well as leaf litter (Zona and Christenhusz 2015). In contrast, the most notable characteristic of TBE is their ability to retain litter and form arboreal soil, although some species do store water in their leaf axils (Derraik 2005, 2009; Derraik and Heath 2005; Killick *et al*. 2014). Also, tank-bromeliads are restricted to the Neotropics, while TBE species inhabit forests from Japan to Africa and southern South America. Moreover, the topic of tank bromeliads has been far more widely studied than TBE: an exploratory search for [“tank bromeliad”] in Google Scholar produces 959 results, while a search for [“trash-basket epiphyte”] produces only 31 citations (February 2021). The lack of attention received by TBE could be related to the fact that arboreal soil is considered as a ubiquitous resource in some forest canopies. However, arboreal soil accumulation and decomposition *in situ* can take a long time due to the low percentage of debris retained by trees (Nadkarni and Matelson 1991). Accordingly, TBE adaptations to enhance litter capture and retention – including their own decaying tissues – may accelerate arboreal soil formation and thus contribute to structuring ecological communities in the canopy by enhancing nutrient availability at the attachment point of each TBE (Hsu *et al*. 2002), ameliorating water and temperature stress (Turner and Foster 2006; Jian *et al*. 2013; Seidl *et al*. 2020), or providing shelter to microbes, invertebrates, and vertebrates that contribute to debris decomposition and nutrient cycles (Figure 1; Paoletti *et al*. 1991; Ellwood and Foster 2004; Karasawa and Hijii 2006b; Ortega-Solís *et al*. 2017; Donald *et al*. 2017; Seidl *et al*. 2020; Donald *et al*. 2020). The influence of TBE on biodiversity and canopy ecosystem functions varies by forest type and scale of analysis. For instance, a few grams of organic matter accumulated in any TBE may not be important for arboreal mammals or large plants but may be crucial for the colonization of mites or bacteria. Yet, we consider that the ecological importance of TBE as secondary FS has been largely neglected in most forest ecosystems.

**Figure 1:**
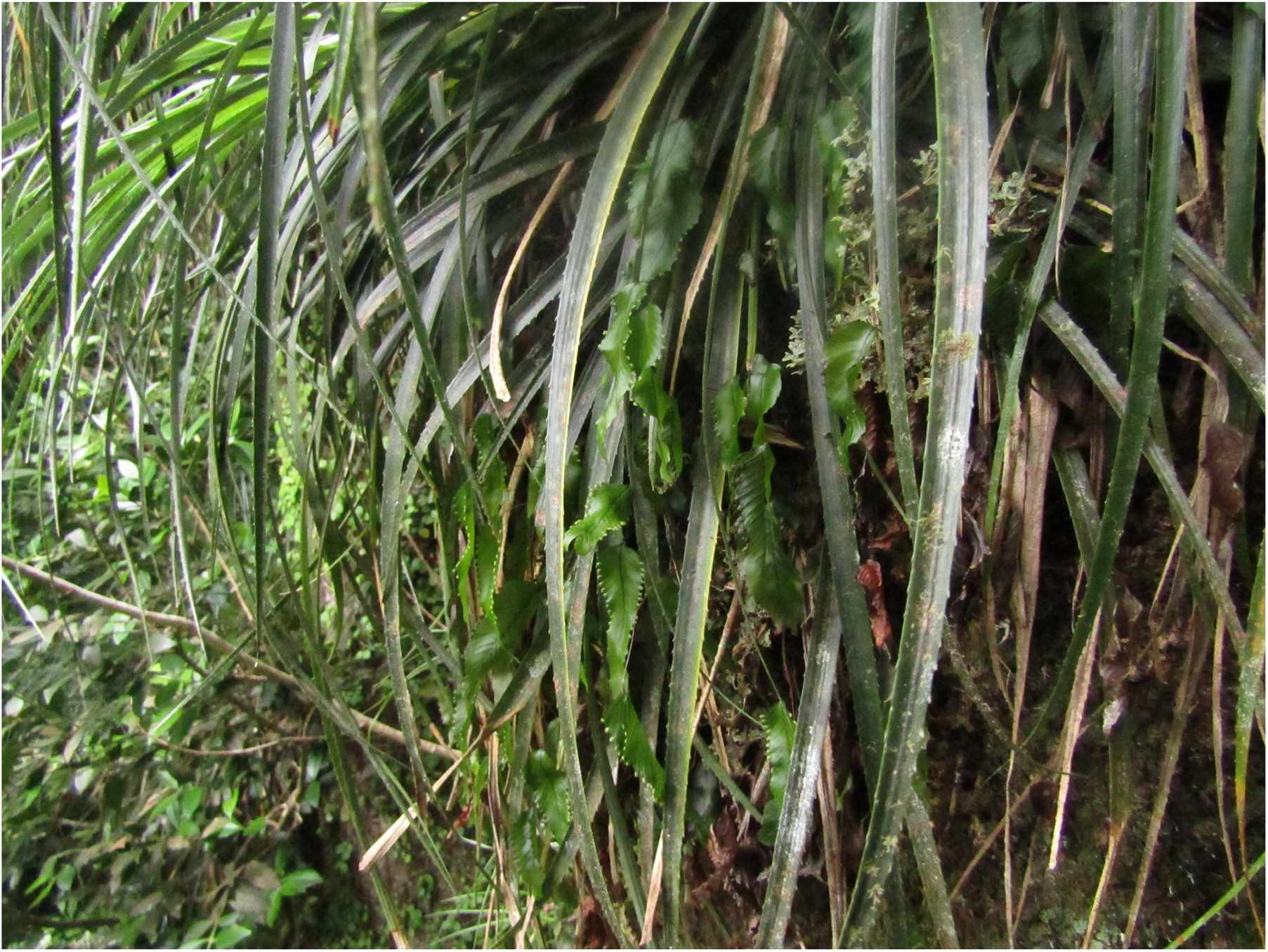
Proposed effects of trash-basket epiphytes on canopy biodiversity and ecosystem functions. Red circles show processes occurring within a TBE, blue circles show animal and plant biodiversity, and the arrows show connections between them. The litter and other debris fall in a TBE **(1)** or decaying leaves and roots are retained within the plant structure **(2)**. Then, nutrients leach from the decomposing organic matter through the stem flow and could reach other epiphytes **(3)**. The decomposition of organic matter trapped by the TBE by bacteria, fungi, and invertebrates **(A)** release nutrients **(B)**, creates a water reservoir **(C)**, and regulates temperature (**D**; high temperatures are lowered, and low temperatures are increased). Other tree dwelling plants (**E**; e.g., epiphytes, accidental epiphytes, lianas, vines) could benefit from the nutrients, water, and temperature regulation provided by TBE **(4,5,6)**, while animals may also benefit from temperature regulation **(7)**. Tree-dwelling plants provide habitat to animals **(8)**, which in turn participate in litter decomposition and nutrient release **(9, 10)**, through litter fragmentation or nutrient deposition from faeces or their carcasses. Images from <u>Openclipart</u>.

Here, we explicitly assess the taxonomic and geographic distribution of TBE species, their participation in ecosystem functioning processes, and their participation in plant-animal and plant-plant interactions in forest canopies worldwide. We applied the TBE concept as defined by Benzing (1990), thus leaving out tank plants, which also act as litter trappers (Zona and Christenhusz 2015). In this review, we address the following questions that are essential for understanding the ecological importance of TBE in forest ecosystems globally: i) How many TBE are presently known worldwide? ii) How many TBE species are present in different forest ecosystems?, iii) How does the presence of TBE influence forest biodiversity and ecosystem functions?, and iv) How can TBE and their plant-plant or plant-animal interactions can be integrated into forest management and conservation efforts? We also highlight unresolved research questions that focus on TBE in forest ecosystems and examine documented interactions of TBE with other trophic groups.

## TAXONOMIC DIVERSITY OF TRASH-BASKET EPIPHYTES

We performed a systematic review of the literature about TBE species. Multiple problems have been acknowledged in published reviews in ecology and environmental sciences, such as lack of a comprehensive and a clear and repeatable selection of records (Haddaway *et al*. 2020). To overcome such pitfalls in our study, we followed the guidelines of the Preferred Reporting Items for Systematic Reviews and Meta-Analyses (PRISMA; Moher *et al*. 2009). Species defined as TBE could also be called humus-collecting plants (Copeland 1907), detritophilic or detritophilous (Cornejo and Iltis 2010), litter-collecting (Lachenaud and Jongkind 2013), litter-trapping (Zona and Christenhusz 2015), litterbin, litter-gathering (Cheek *et al*. 2008; Lachenaud and Jongkind 2013), humilectic plants (Dressler 1981), bird’s nest ferns, nest-epiphytes (Schimper and Fisher 1903), and litter-impounding plants (Zotz 2016). Thus, we collected documents about TBE in Google Scholar, Google Books, Microsoft Academic, and Web of Science using search operators to combine all the names a TBE could receive in a single string: [“epiphyte” AND (trash-basket OR trashbasket OR cruddophyte OR detritophil OR humilectic OR humus-collect OR litter-collect OR litter-gather OR litter-trap OR litterbin OR nest fern OR nest-epiphyte OR litter-impound)]. We used the option “Search in all databases” in Web of Science and performed a search in titles and abstracts. Microsoft Academic is a semantic search engine that interprets queries differently than Google’s search engines, so we had to use the tool “Top topics” to restrict the results to the topic Ecology. Our Google Books search was restricted to records with an available preview, which allowed us to search the book content. After an initial examination of the collected records, we retrieved additional references cited in the literature. We used Zotero (Stillman *et al*. 2020) for record completion, screening, classification, and merging duplicate entries. After the first screening of TBE species, we repeated the search in Web of Science, Google Scholar, and Google Books with the genus and the string [invertebrate OR plant-animal OR plant-plant] to find more records describing interactions between TBE and other organisms. We also searched the scientific names and synonyms of TBE species in Web of Science to record the number of articles per species. In this way, we were able to identify the TBE that lacked attention from researchers. We compared the count of articles per species returned by Web of Science with the yearly distribution of the 120 studies supporting the inclusion of a species in our list of TBE. We summarised the records collection and their processing in a PRISMA flow chart (Supplementary Information Figure 5).

We compiled a global list of 209 TBE species from 120 records (57 research articles, 39 reviews, 8 books, 6 book chapters, 7 theses, one comment, one proceeding, and one technical report), which provided pictures, morphological descriptions, personal observations of the authors, or sampled the litter or soil trapped by an epiphyte. The taxonomic information of each TBE species was standardized according to World Flora Online (WFO 2021). A set of 30 TBE species appear as ambiguous in WFO (they have not been assigned to any accepted names as of February 2021) and 4 were not found but are recognized as valid species in Plants of the World Online (POWO 2019), so we kept them in our database but marked them as ambiguous, since further updates to WFO could result in accepting their names or assigning them to other species. The epiphytic life form of most TBE was verified in Epilist 1.0 (Zotz *et al*. 2021) and Plants of the World Online (POWO 2019). However, the strict assignment of a life form was difficult in some cases, and we included in our database 10 species that could not be unequivocally designated as epiphytes, but that share the litter-trapping function with the rest of the species on the list.

TBE appear to have evolved many times independently within a wide range of plant families. Of 209 TBE species reported in the literature, 43% belong to Araceae, 23% to Polypodiaceae, 14% to Orchidaceae, 9% to Aspleniaceae, and 11% distributed among Asteliaceae, Bromeliaceae, Cyclanthaceae, Dryopteridaceae, and Pandanaceae (Tables 1 and 2, and Figure 2). Most reports focus on Aspleniaceae and Polypodiaceae with 36% and 26% respectively, half of all TBE species (Table 2). However, we found most of the TBE species in the Araceae are probably due to the large number of *Anthurium* species inferred from the taxonomic descriptions in Croat (1991; 64 species, see Supplementary Information). TBE in the Polypodiaceae and Aspleniaceae have attracted the most research attention according to the number of records in Web of Science. At the same time, 93 species were not mentioned in any of the literature databases gathered by Web of Science. The families with the highest number of TBE species lacking studies are Araceae (65 species) and Polypodiaceae (18 species). The lack of studies in Araceae also explain the concentration of non-studied species within the Neotropical realm (see Supplementary information). The studies about TBE found date back to 1904 and started to appear regularly during the last 50 years. General research interest in the species listed here as TBE increased faster than studies about their role as TBE. The lack of studies about 93 TBE species does not imply that they are completely unknown because Web of Science does not index the full text of articles, and the species could be covered in theses or the grey literature. However, this result strongly suggests that some species receive less attention in the main research outlets.

**Figure 2:**
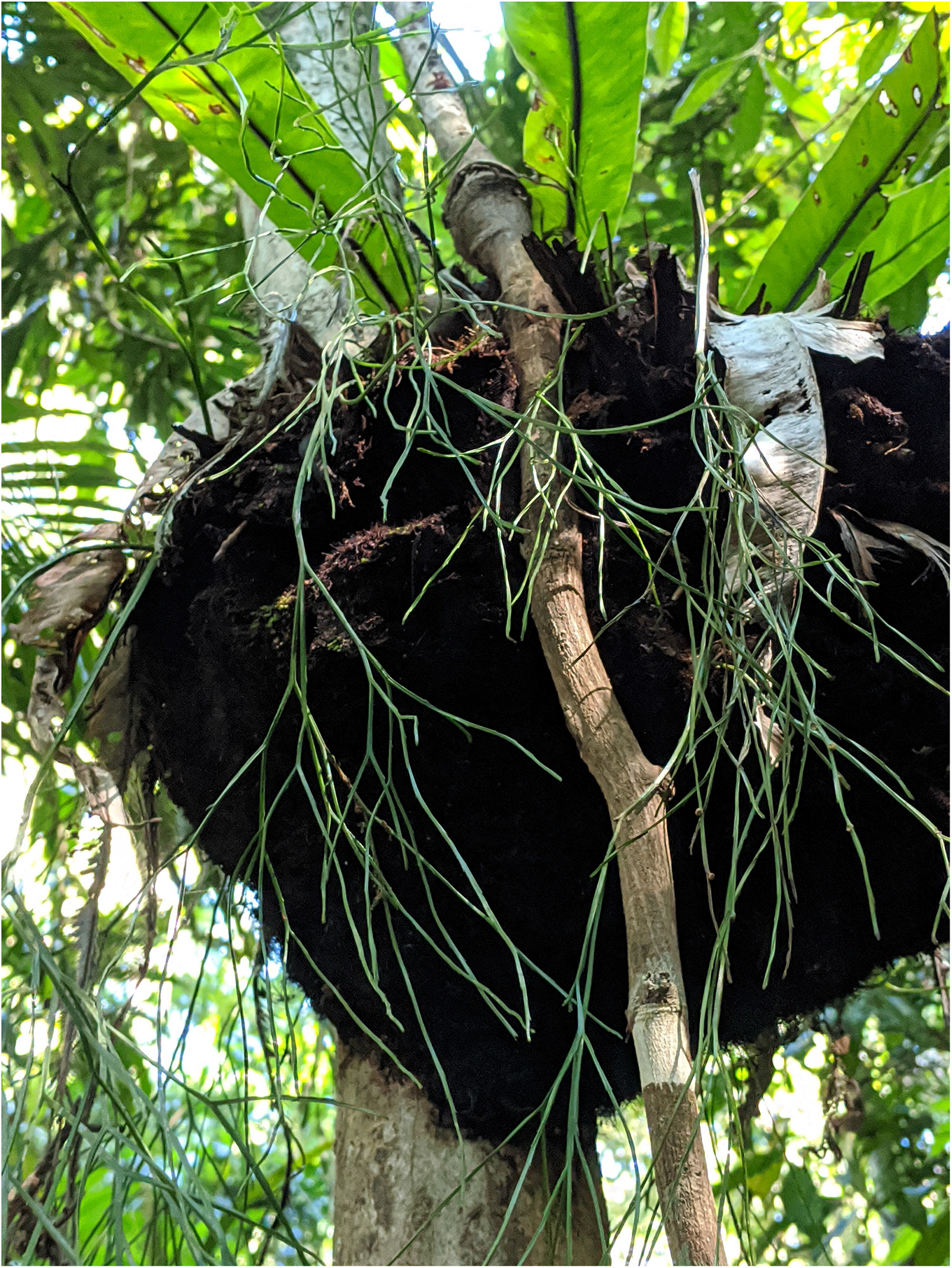
Examples of trash-basket epiphytes. **A:** Litter and other debris trapped by *Platycerium bifurcatum*, Danbulla National Park, Australia. **B:** Clump of *Fascicularia bicolor* in *Eucryphia cordifolia* tree, Chiloé Island, Chile. **C:** Fertile (green) and sterile (brown) fronds of *Drynaria rigidula*, Danbulla National Park, Australia. **D:** A close-up of *Aglaomorpha heraclea* showing fronds widened at the base, Philipines. Credits: H. Brent (A and C), D. Mellado-Mansilla (B), and M. Kessler (D).

**Table 1:** Distribution of TBE per family, number of records supporting species designations as TBE, and the percentage represented by each family. Data are based on 108 publications that described a species using the term TBE or related concepts, had pictures that show litter trapped by an epiphyte, or sampled the litter and soil of a TBE species. The columns **Species lacking studies and Percentage without studies per family** refer to the number of TBE species without studies recorded in the Web of Science, relative to the total number of TBE identified within each family. *The number of references sums more than 108 because some of them provide information on several families.

| Family | TBE Species (%) | References (%)* | Species lacking studies | Percentage without studies per family |
| --- | --- | --- | --- | --- |
| Orchidaceae | 29 (14) | 16 (11) | 6 | 21% |
| Polypodiaceae | 49 (23) | 39 (26) | 18 | 37% |
| Araceae | 90 (42) | 16 (11) | 65 | 72% |
| Aspleniaceae | 19 (9) | 53 (35) | 6 | 32% |
| Cyclanthaceae | 2 (1) | 2 (1) | 0 | 0% |
| Asteliaceae | 4 (2) | 10 (7) | 0 | 0% |
| Dryopteridaceae | 17 (8) | 12 (8) | 4 | 24% |
| Bromeliaceae | 1 (<1) | 1 (1) | 0 | 0% |
| Pandanaceae | 2 (1) | 1 (1) | 2 | 100% |

**Table 2:**
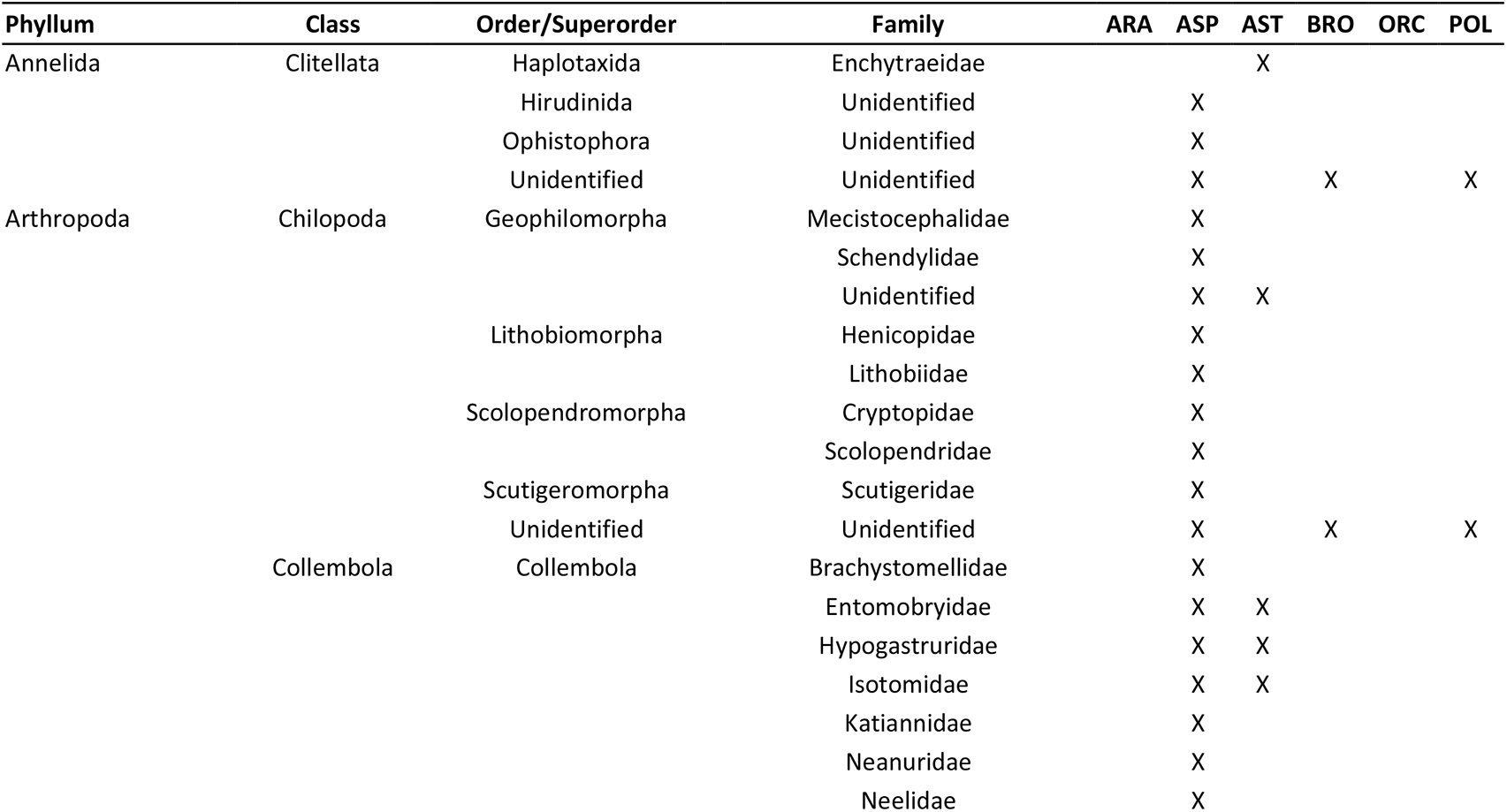
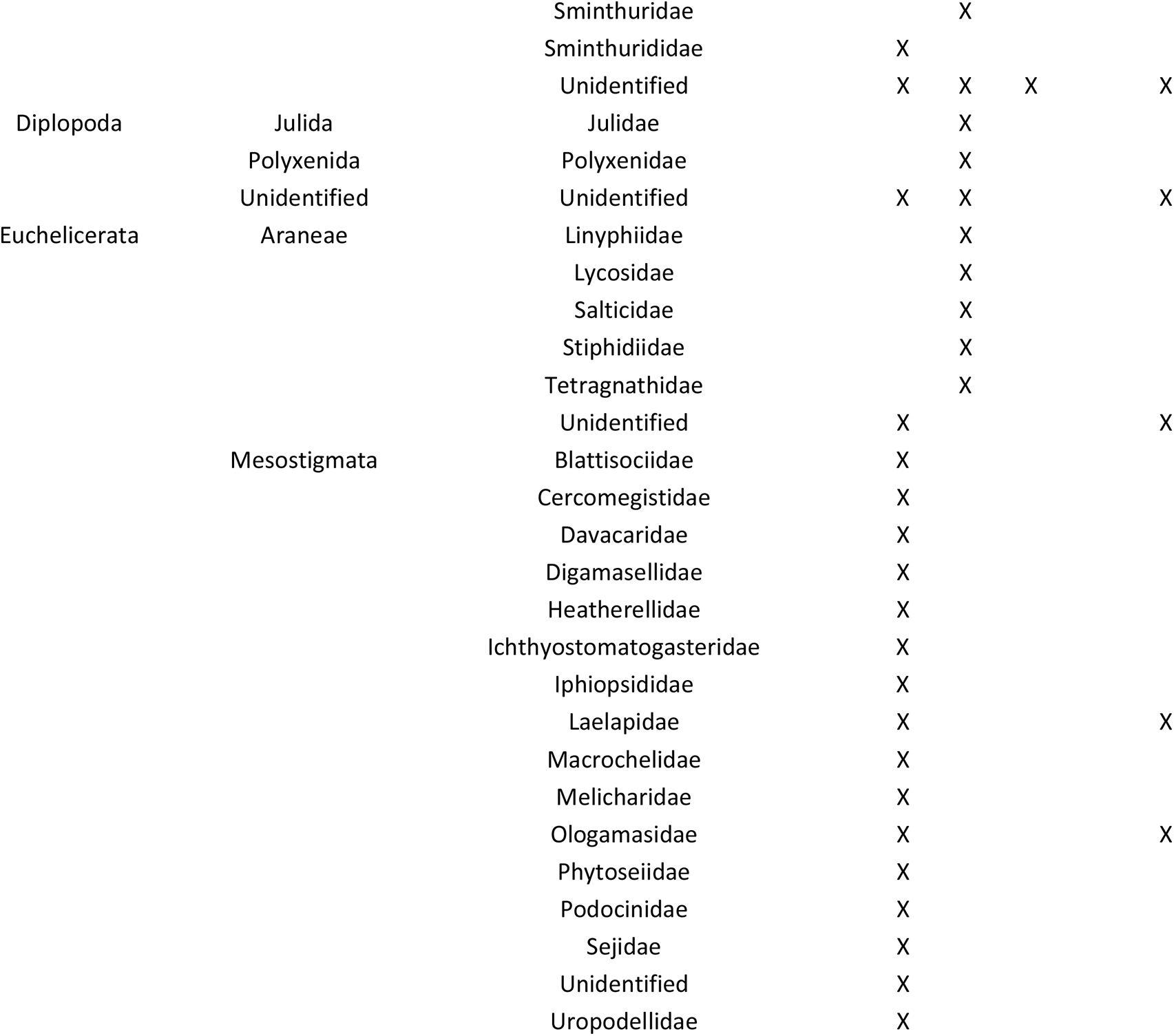
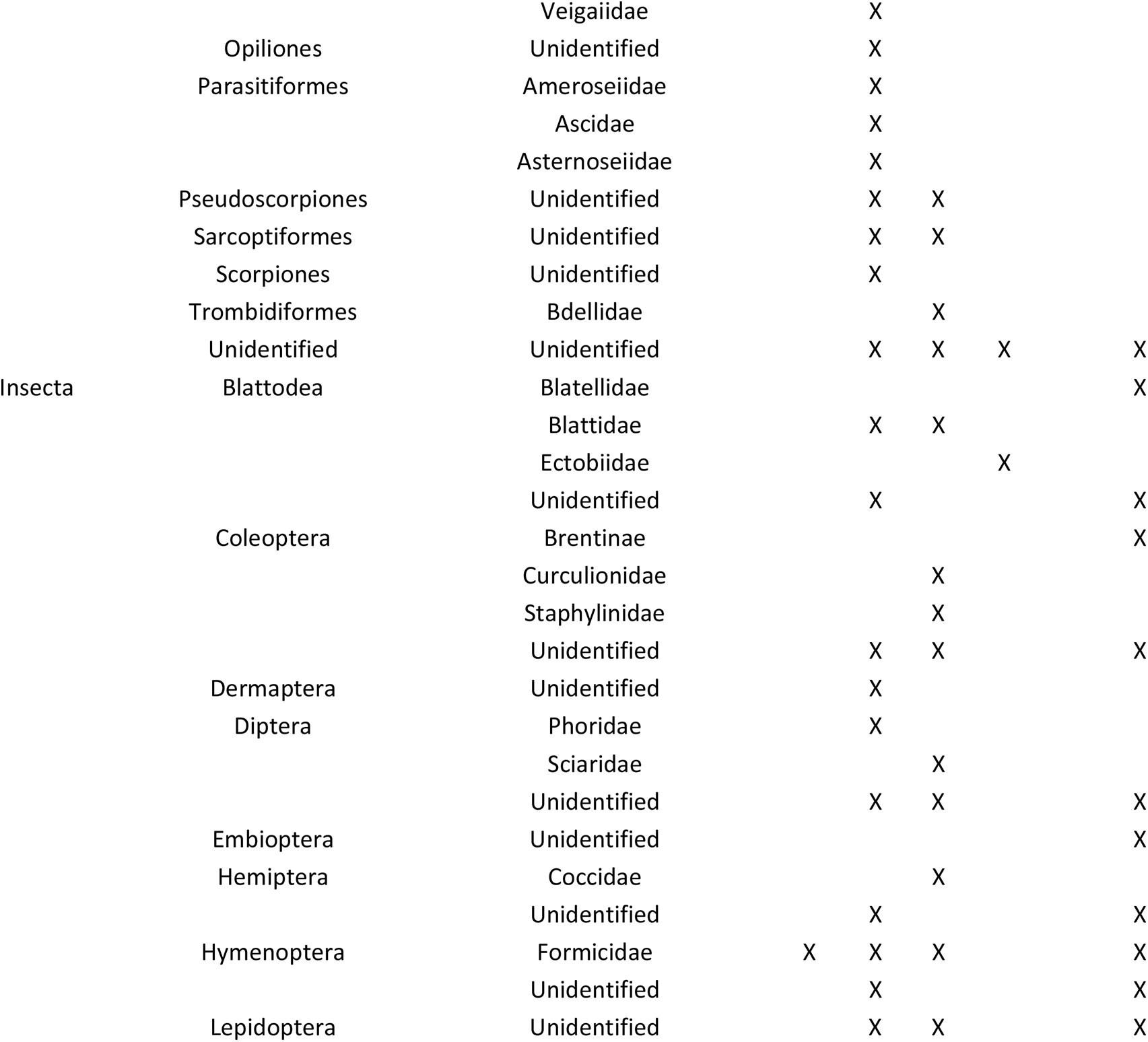
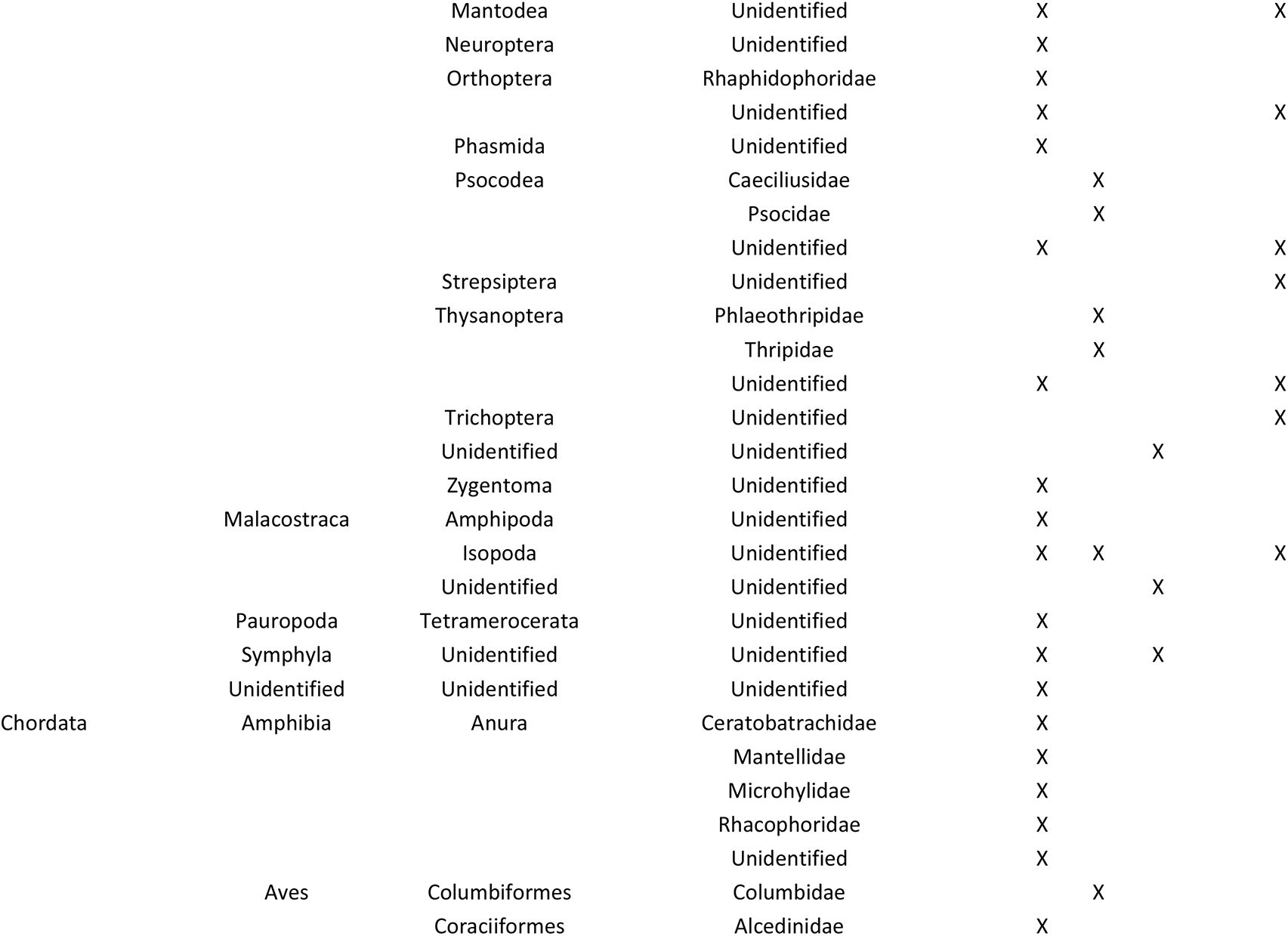
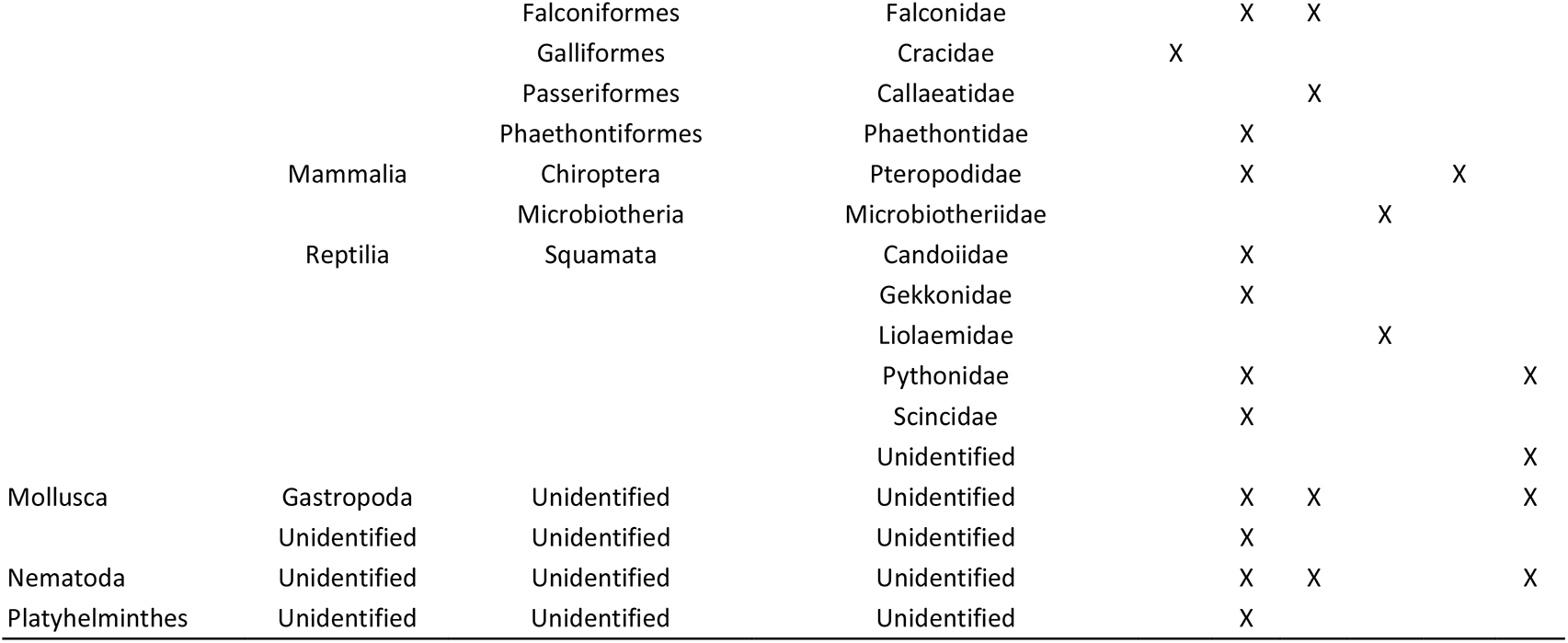
Animal families (and higher categories) reported in the literature to interact with TBE family globally. Unreported taxonomic levels were filled as “Unidentified”. **ARA**: Araceae, **ASP**: Aspleniaceae, **AST**: Asteliaceae, **BRO**: Bromeliaceae, **ORC**: Orchidaceae, **POL**: Polypodiaceae. A detailed list of references is provided in Supplementary Information.

A comparison of the number of species identified as TBE and the number of epiphytes per family suggests that additional TBE species could potentially exist, specially within speciose epiphytic families. For instance, 34 orchid species from 15 genera have been described as TBE, from 559 genera and ca. 21,169 species of epiphytic orchids (Zotz *et al*. 2021). However, the orchid genera that include TBE species amount to 4,097 epiphytic species (Zotz *et al*. 2021), meaning that less than 1% of them are in our list. There are striking differences in the genera *Bulbophyllum* and *Dendrobium* because only one TBE has been reported in each of them, yet they contain 2,043 and 1,557 epiphytic species, respectively (Zotz *et al*. 2021). More TBE could also occur in Aspleniaceae or Dryopteridaceae. We identified 19 TBE within 246 known epiphytic species of *Asplenium* (Zotz *et al*. 2021). However, “nest ferns” mostly occur in section **Thamnopteris** Presl., suggesting that the number of non-reported TBE in Aspleniaceae could be small. Regarding Dryopteridaceae, 17 TBE have been described within the genus *Elaphoglossum*, which includes 415 epiphytes (Table 1). While we cannot assume that there is a high number of TBE that remain unreported within the families having the largest number of epiphytic species, we suggest that more species could be described in future studies, especially within epiphytic orchids. On the other hand, Polypodiaceae seem well represented among TBE with ca. 30% of the known epiphytic species listed (Table 1).

Whether litter-trapping structures were involved in the evolutionary transitions to the epiphytic habit remains unclear. Nevertheless, this trait has been suggested as a driver of the diversification in the epiphytic Drynarioid ferns (Janssen and Schneider 2005). On the other hand, nest ferns within Polypodiaceae have been shown to diversify at a slow rate – lower to non-differentiable from the background diversification rate of the family (Sundue *et al*. 2015). TBE attributes could have been already present in terrestrial ancestors or could have evolved within epiphytic species to compensate for the lack of soil substrate in the canopy. Although species diversification can be driven by multiple factors (Sundue *et al*. 2015), it could be interesting to evaluate whether diversification rates among multiple epiphytic taxa are correlated with the ability of species to “recreate” a terrestrial habitat above trees. We suggest that studies about the taxonomy and evolution of epiphytic clades should assess litter-trapping and the consequent habitat formation among the potential factors affecting species diversification.

## GEOGRAPHIC DISTRIBUTION OF TRASH-BASKET EPIPHYTES

We calculated the total species richness per forest biome and ecoregion by cross-referencing the native ranges of 201 TBE found in Plants of the World Online database (POWO 2019) with spatial polygons of terrestrial ecoregions (Olson *et al*. 2001; Figure 3). Additionally, we assessed the effect of biomes on the diversity of TBE using a generalized linear model (GLM) with a Poisson distribution. Because species richness varies with area, we constructed 8 grids with hexagonal cells and areas ranging from 12,000 to 774,400 square kilometres. The area of some cells was reduced after cross-referencing the grids with landmasses; therefore, we kept the cells larger than 12,000 square kilometres and with at least 1 TBE occurrence. Since cells could overlap with more than one biome, we used the percentage cover of each biome per cell as predictor variables in our GLM. We fitted two GLMs, one excluding the Tropical & Subtropical Moist Broadleaf Forests (TSMBF) and the other excluding the Tropical & Subtropical Grasslands, Savannas & Shrublands (TSGSS) because their percentage cover were negatively correlated (r = -0.6) producing variance inflation. The total richness of vascular plants per ecoregion could influence the number of TBE per cell, thus, we used the data in Kier *et al*. (2005) to create a ranking of total plant richness per ecoregion and added the corresponding value to each cell. We estimated a weighted mean of the plant richness rank for cells containing more than one ecoregion. The cell area and plant richness rank were scaled around their median and median absolute deviations. After adding the corresponding data to each cell, we pooled the grids, took a random sample of 500 cells without replacement, and fit an additive GLM with the form:

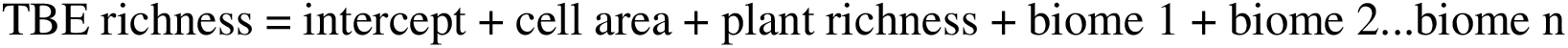

**Figure 3:**
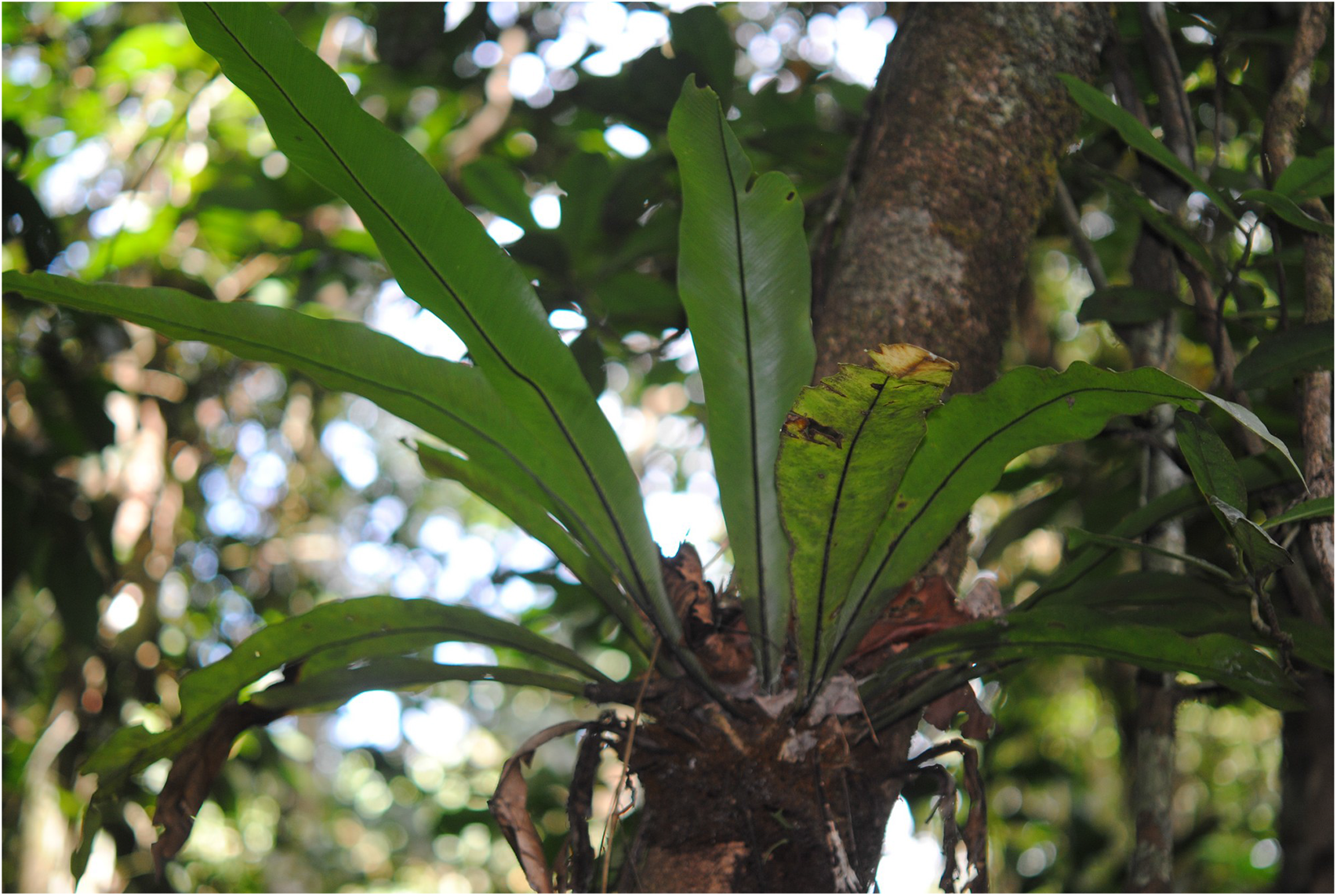
Geographic distribution of 201 TBE species globally. **Panels A** and **B** show the TBE species richness per terrestrial ecoregion and biome. **Panel C** shows the estimated coefficients of Poisson model of TBE richness. **TSMBF**: Tropical & Subtropical Moist Broadleaf Forests, **TSDBF**: Tropical & Subtropical Dry Broadleaf Forests, **TSGSS:** Tropical & Subtropical Grasslands, Savannas & Shrublands, **Mngr**: Mangroves, **TSCF**: Tropical & Subtropical Coniferous Forests, **TBMF**: Temperate Broadleaf & Mixed Forests, **FGS**: Flooded Grasslands & Savannas, **TmCF**: Temperate Conifer Forests, **TGSS**: Temperate Grasslands, Savannas & Shrublands, and **MFWS**: Mediterranean Forests, Woodlands & Scrub (Olson et al. 2001).

The maximum distance required to ensure that all cells in a random sample had at least one neighbouring cell was 3,244 km (estimated from 1,000 samples of 500 cells). Therefore, we performed a permutational Moran’s I test for distance bands ranging from 110 to 6,488 km and found spatial dependence on the residuals at all distances (p < 0.01). We re-fit the non-spatial GLM adding a set of Moran eigenvectors to the model formula to account for the spatial auto-correlation of the residuals. The Moran eigenvector approach consists of finding one or more numeric vectors that fit part of the spatial noise in the model (Griffith and Peres-Neto 2006). We tested different distance bands to generate the Moran eigenvectors and selected the one corresponding to 4,610 km of distance (Moran’s probability > 0.05). After checking model assumptions, we applied the final model to 1,000 samples and averaged the parameter estimates across all the fitted models. Model averaging allowed us to reduce the influence of the spatial distribution of each sample on the parameters estimated in our GLM. All analyses were performed in R 4.0 (R Core Team 2020) with the packages spocc (Chamberlain *et al*. 2018), taxize (Chamberlain and Szöcs 2013), spatialreg (Bivand *et al*. 2013), and MuMIn (Bartoń 2020). Additional details of the methods are provided as R scripts in Supplementary Information.

TBE are widely distributed across four continents, with maximal species richness occurring between latitudes 20° and -20° (Figure 3). The forest biomes with the highest total richness of TBE are Tropical & Subtropical Moist Broadleaf Forests (TSMBF; 184 species), Mangroves (159), and Tropical & Subtropical Dry Broadleaf Forests (TSDBF; 147 species). Within ecoregions, the South American Pacific mangroves (80 species), Eastern Cordillera Real montane forests (67 species), and the Napo moist forests (67 species) had the highest number of reported TBE. These three ecoregions with the highest species richness of TBE occur in the Neotropical realm and are located in Colombia, Ecuador, and Perú, respectively, with the South American Pacific mangroves extending also to Panamá. As expected, our spatial GLM shows that the number of TBE species increases with the total plant species richness or the area of each grid cell. An increase in the cell coverage of TSMBF had positive effects on the number of TBE while other biomes have negative or neutral effects (Figure 3). The peak of TBE species richness occurs in centres of high vascular plant diversity within the TSMBF (Figure 3). The lowest species richness of TBE in Sub-Saharan Africa (Figure 3) coincides with Africa’s low species richness of plants compared to forests in the same latitude (Kier *et al*. 2005). Moreover, large epiphytic families such as Araceae and Orchidaceae – which contain 57% of species in our dataset – are underrepresented in the Afrotropics (Taylor *et al*. 2021). In summary, the geographical distribution of TBE follows the same patterns as those of the species richness of vascular plants, which is primarily associated with higher potential evapotranspiration, number of wet days per year, and habitat heterogeneity (Kreft and Jetz 2007). The TSMBF is consistently the most important forest biome for TBE, even when controlling for the effect of its larger area (Figure 3). The TSMBF extends across the Neotropical, Afrotropical, Indomalayan, and Australasian realms, thus, comprising a mixture of floras of different evolutionary origins that increases the probability of having multiple TBE species. On the other hand, the lower TBE species richness in Mediterranean Forests, Woodlands & Scrub, Temperate Broadleaf & Mixed Forests, Temperate Grasslands, Savannas & Shrublands, and the Tropical & Subtropical Grasslands, Savannas & Shrublands reflects the lower representation of epiphytes within these biomes. Mangroves show a strikingly wide confidence interval in our spatial model (Figure 3). Thus, despite being among the biomes with high TBE species richness, their importance requires further evaluation. Mangroves are the smallest biome, covering only 346,436 km^2^, thus, we did not have any cells that were completely covered by mangrove. However, Mangroves often occurred in cells dominated by TSMBF in our dataset, and often covered small portions of cells with more than 20 TBE. Moreover, 100% of TBE inhabiting Mangroves are shared with the TSMBF (Supplementary Information). Therefore, in some TBE-rich areas, mangroves could represent extensions of their neighbouring biomes.

We emphasize that biogeographical analysis reported here may be subject to well-known taxonomic and geographic biases (Hortal *et al*. 2015). The number of TBE could be larger in understudied taxa or under-studied ecosystems. As stated before, because trash-basket orchids could be under-reported in the literature, this could lead to changes in the richness trends reported here. Another key consideration is that TBE is a sub-group within a larger group of epiphytic litter-trappers (Zona and Christenhusz 2015). Consequently, in some ecoregions, the same function could be performed by epiphytes that are not included in the strict definition of TBE, such as Neotropical tank bromeliads. Tank bromeliads can be found in 25 plant genera (Zona and Christenhusz 2015) that include more than 1,500 epiphytic species (Zotz et al. 2021), but that only occur in the Americas, from the US to northern Argentina. Therefore, if they are studied together with TBE it is likely that tank bromeliads will increase the importance of litter trapping in Neotropical ecoregions.

## LITTER-TRAPPING STRATEGIES AND FUNCTIONAL CONSEQUENCES

TBE display multiple litter trapping strategies even within the different taxonomic groups. *Asplenium* and *Microsorum* nest ferns have a rosette shape with leaves oriented upwards, making a funnel that channels the litter and debris towards the stem (Zona and Christenhusz 2015). Within Polypodiaceae, the genus *Platycerium* has sterile fronds that cover the roots and form an open crown where organic matter and water accumulate (Dubuisson *et al*. 2009; Figure 2A). *Drynaria* and *Aglaomorpha* ferns have adaptations that range from normal leaves widened at the base (Figure 2D) to completely dimorphic leaves with specialized litter-trapping leaves that are short, stiff, and dead yet remain attached to the fern (Janssen and Schneider 2005; Dubuisson *et al*. 2009; Vasco *et al*. 2013; Figure 2C). Araceae epiphytes have a rosette shape, but they also form a trash-basket with specialized roots pointing upwards (Zona and Christenhusz 2015). The same type of trash-basket roots seems to be the most common litter-trapping mechanism in Orchidaceae, but litter trapping in this family requires more studies to quantify the number of species and the range of litter-trapping strategies they use. Asteliaceae and the bromeliad genus *Fascicularia* form clumps, which create an intricate surface where litter is trapped (Figure 2B). The spines along the margins of *Fascicularia* leaves could be also helpful to reduce the loss of litter by trapping leaves that are blown away by the wind (G. Ortega-Solís, obs. pers.).

The contribution of TBE to arboreal soil accumulation in a forest stand could be decoupled at two scales: 1) the microhabitat – represented by the litter holding capacity of each single individual – and 2) the forest stand scale that considers the relative contribution of TBE to the overall accumulation of arboreal soil in the forest. At the microhabitat scale for example, the dry mass of litter and arboreal soil held by a single nest fern could range between 10 to 18% of its fresh weight (Pócs 1980). The fresh weight of nest ferns filled with organic material could vary from 1 to 200 kg per individual (Fayle *et al*. 2009), therefore we can roughly infer that each nest fern could hold from 0.1 to 20 kg of dry mass of litter, other debris, and arboreal soil. At the forest stand scale, the relative contribution of TBE to arboreal soil accumulation in different forest ecosystems should depend on the abundance of TBE species and individuals, and the successional stage of the forest. For instance, the litter and arboreal soil of nest ferns represent 8.6% (0.18 Mg ha^-1^) of the dry matter of epiphytic material (biomass+arboreal soil) in sub-montane forests of Tanzania (Pócs 1980) and 10% (0.3 Mg ha^-1^) in the subtropical forests from Taiwan (Hsu *et al*. 2002), but represents approximately 100% of the arboreal soil both studies report. The abundance of *Asplenium nidus* and *Asplenium phyllitidis* in the canopy of lowland rainforests from Borneo ranges from 0-146 and 0-414 ind/ha respectively (Fayle *et al*. 2009). *A. nidus* is more abundant in forest stands with a higher occurrence of emergent trees and an open understory, while *A. phyllitidis* increases in stands with closed canopy (Fayle *et al*. 2009) so their contribution to arboreal soil accumulation varies depending on the same attributes that determine their abundances. In South American temperate rainforests, *Fascicularia bicolor* contributes between 27 and 57% of the total dry matter of the arboreal soil recorded in a single host-tree species (Díaz *et al*. 2010). *F. bicolor* reaches densities between 575 and 1,675 ind/ha (G. Ortega-Solís, unpublished data); thus its contribution of arboreal soil may vary widely depending on the availability of large host-trees (Ortega-Solís *et al*. 2020). We highlight that most of the data available about the relative importance of the arboreal soil held by TBE is biased towards large TBE species in forests where their contribution is significant. Future studies of canopy soils and epiphytic biomass should pay more attention to the separate contributions of all species of epiphytes in forests. Although this will be logistically difficult, it will help clarify the relative importance of single species or functional groups of epiphytes for ecosystem functions associated with arboreal soils, such as nutrient cycling and water retention, and how such functions change across forest successional age or under different types of forest management.

The litter and soil of TBE are sources of nutrients for themselves and associated organisms. The nitrogen content in nest ferns’ soil is like that found in non-TBE arboreal soils. For instance, N concentrations of 23.1 mg g^-1^ (sd = 3) and 18.7 mg g^-1^ (sd = 2.3) have been reported for arboreal soils within and outside TBE, respectively (Hsu *et al*. 2002). The leaves of epiphytes rooted in arboreal soil have a higher content of nitrogen than epiphytes lacking soil (Hietz *et al*. 2002). Because litter is a primary source of nitrogen, which is a limiting nutrient in many forests, the litter retention capacity of TBE could allow them and their associated plants to be less dependent on atmospheric inputs of nitrogen through symbiotic or diazotrophic N-fixers. However, an increase in nitrogen availability can also lead to changes in the relative availability of other nutrients. For instance, the phosphorus content in TBE soil is a quarter of the concentration found in other arboreal soils in Taiwan forests, leading to an N:P ratio of 100:1 in TBE soil, compared to 23:1 in arboreal soils not associated with TBE (Hsu *et al*. 2002). Accordingly, phosphorus (or other macro- and micronutrients; John *et al*. 2007) could be more limiting than nitrogen for TBE and plants growing on their associated arboreal soils (Huang and Lin 2016). Studies on competition between plants could provide valuable information on how TBEs interact with their host trees and the plants colonizing TBE’s soils. There could be competition for nutrients – and a possible deprivation of nutrients for the TBE – or the colonization by other epiphytes could increase the pool of available nutrients by enhancing litter retention. In addition, future studies may also focus on the effects of TBE on their host trees. On the one hand, the increased provision of nutrients could be favourable to host trees with adventitious roots that take up nutrients from the arboreal soils (Nadkarni 1981), while on the other hand, the weight associated with large TBE individuals could increase the risk of tree or branch fall.

While TBE create a nutrient reservoir that can be used by themselves and by other organisms, they could also reduce nutrient availability for plants living underneath (Chen *et al*. 2019). For example, *A. nidus* has been observed to decrease the concentrations of potassium and magnesium in stem flow (Chen *et al*. 2019). This reduction could be related to the high nutrient demand of TBE, but also could be a dilution effect because the leaves of *A. nidus* channel throughfall toward the stem (Chen *et al*. 2019). Regarding nitrate concentration measured in the stem-flow above and below nest ferns, responses vary from no effect (Chen *et al*. 2019) to an increase of 85% (Turner *et al*. 2007). An increase in nitrate concentration could be expected because of litter decomposition within TBE. Yet, a potential explanation for lower nitrate concentrations in stem flow could be due to the acid soil pH in TBE that reduces the presence of nitrifying bacteria (Vance and Nadkarni 1990), hence promoting less water-soluble forms of nitrogen and reducing the leaching losses. It is likely that TBE increase nutrient availability at their point of attachment to the host trees (acting as secondary FS at the microhabitat scale), thus creating a flow of nutrients between the TBE, its soil, and organisms inhabiting in it (Figure 1). However, in the long term, nutrients will inevitably be released and incorporated into the larger nutrient cycle of the forest when the TBE senesces or is dislodged by extrinsic forces.

TBE’s soil contributes to water retention in the canopy and consequently influences air temperature and humidity in the space of a few centimetres around tree branches (Freiberg 1996, 2001). The extent of the influence of TBE on water retention and microclimate in the forest canopy depends on their relative importance within the total epiphytic material in a forest stand and the study length. For instance, arboreal soils in nest ferns represent 3% (497 L ha^-1^) of the water retention capacity of suspended soils in sub-montane forests from Tanzania, while bryophytes and filmy ferns are responsible for as much as 90% water retention (Pócs 1980). However, bryophytes have been reported to double water loss by evapotranspiration compared to canopy humus (Köhler *et al*. 2007), which suggests that large accumulations of arboreal soil – such as the soil in some TBE – could keep water retained for a longer time than bryophytes. The shading effect of fern leaves may also be an important factor in determining long water retention times in arboreal soil, as the weight loss of the root-ball of *A. antiquum* subject to artificial drought is three times faster when the fronds are clipped (Jian *et al*. 2013). The water stored in TBE soil should also ameliorate temperature and humidity variation (Freiberg 2001). In the tropical forests of the Guyanas, the temperature measured 1 cm away from the surface of branches covered by epiphytic material is more than 2°C lower than that of bare branches during the hottest hours of the day (Freiberg 2001). The buffering effect of epiphytic material on air temperature decreases rapidly, but it could be still ca. 1°C at 12 cm distance (Freiberg 2001). Since water is also important for decomposition, it indirectly heats the air around branches producing more than 0.5 °C warmer temperatures than bare branches at night (Freiberg 2001). The temperature regulation by TBEs also depends on the dominant microclimate in each forest. For instance, temperature measurements in bird’s nest ferns in Madagascar were 4.4°C colder than the daily maximum recorded under the same forest canopy, but there were no differences when comparing the minimum temperature in ferns with the daily minimum (Seidl *et al*. 2020). It is likely that in the warm conditions of Madagascar, the temperature at night is not low enough to be different from that measured in the TBE. Even in oil palm plantations, nest fern TBE regulate air temperature and humidity to a similar extent as in native forests (Fayle *et al*. 2010).

## INFLUENCE OF TRASH-BASKET EPIPHYTES ON FOREST BIODIVERSITY

We compiled a database of 48 articles describing plant-animal interactions and 26 describing plant-plant interactions involving a TBE species. The records were collected from Google Scholar, Google Books, Microsoft Academic, and Web of Science during the search for records supporting our list of TBE species (see section “Taxonomic diversity of trash-basket epiphytes”). We considered an article as valid for inclusion in our database of interactions when the interaction described was clearly mediated by the litter-trapping function of a TBE, thus, leaving out pollination. Field observations about two additional animal families and one plant species were added to the data of species interactions. Taxonomic information was standardized according to the Integrated Taxonomic Information System (ITIS; www.itis.gov) for animals and the World Flora Online (WFO; www.worldfloraonline.org) for plants. We report the minimum number of animal families interacting with TBE due to taxonomic uncertainties in some studies. Unfortunately, most articles did not provide enough information to conduct a formal meta-analysis, especially those describing plant-plant interactions. Therefore, we independently summarised the results of two studies comparing the diversity of soil invertebrates between a TBE species and its host tree. We extracted 53 pairs of interactions between TBE and invertebrate families that inhabited them and calculated the average species richness or abundance per sample for each study. Then, we estimated the mean Relative Interaction Index (RII; Armas *et al*. *A.* 2004) for each study and its bootstrapped confidence intervals (BCa method). The RII is a standardized measure of species interactions designed to be symmetric around zero and restricted between -1 and 1. Therefore, the RII is useful to quantify facilitation or competition among a group of interacting species, where RII values close to 1 represent positive effects (facilitation) and -1 negative effects (competition).

TBE structures are inhabited by animals from 97 families belonging to 15 classes (Table 2), including the description of new species (Disney 1999; Vera and Schapheer 2018). Most of the reported animals are invertebrates that inhabit litter and soil (90 families). However, TBE structures are also used by amphibians (4 families), reptiles (5 families), birds (6 families), and mammals (2 families; Table 2). Most faunal reports for TBE are concentrated in the Aspleniaceae (65% of the records). Two studies comparing the richness of soil invertebrates found on TBE and their host-trees showed RII values of 0.30 and 0.38, but confidence intervals overlapped zero in both cases. However, the RII of soil invertebrate abundances shows significant positive effects with means of 0.41 and 0.76 (Figure 4). An increase in species abundance but not richness could be due to the fact that TBE species increased the amount of microhabitat within their host trees, rather than providing new habitats different from those already available on the tree (Figure 5; see also Angelini and Silliman 2014). Regarding plants, at least 72 species from 35 families have been found in TBE (Table 3). The dominant families of plant species that interact with TBE are Hymenophyllaceae and Orchidaceae (11% species each), and Aspleniaceae (10%) (Table 3).

**Figure 4:**
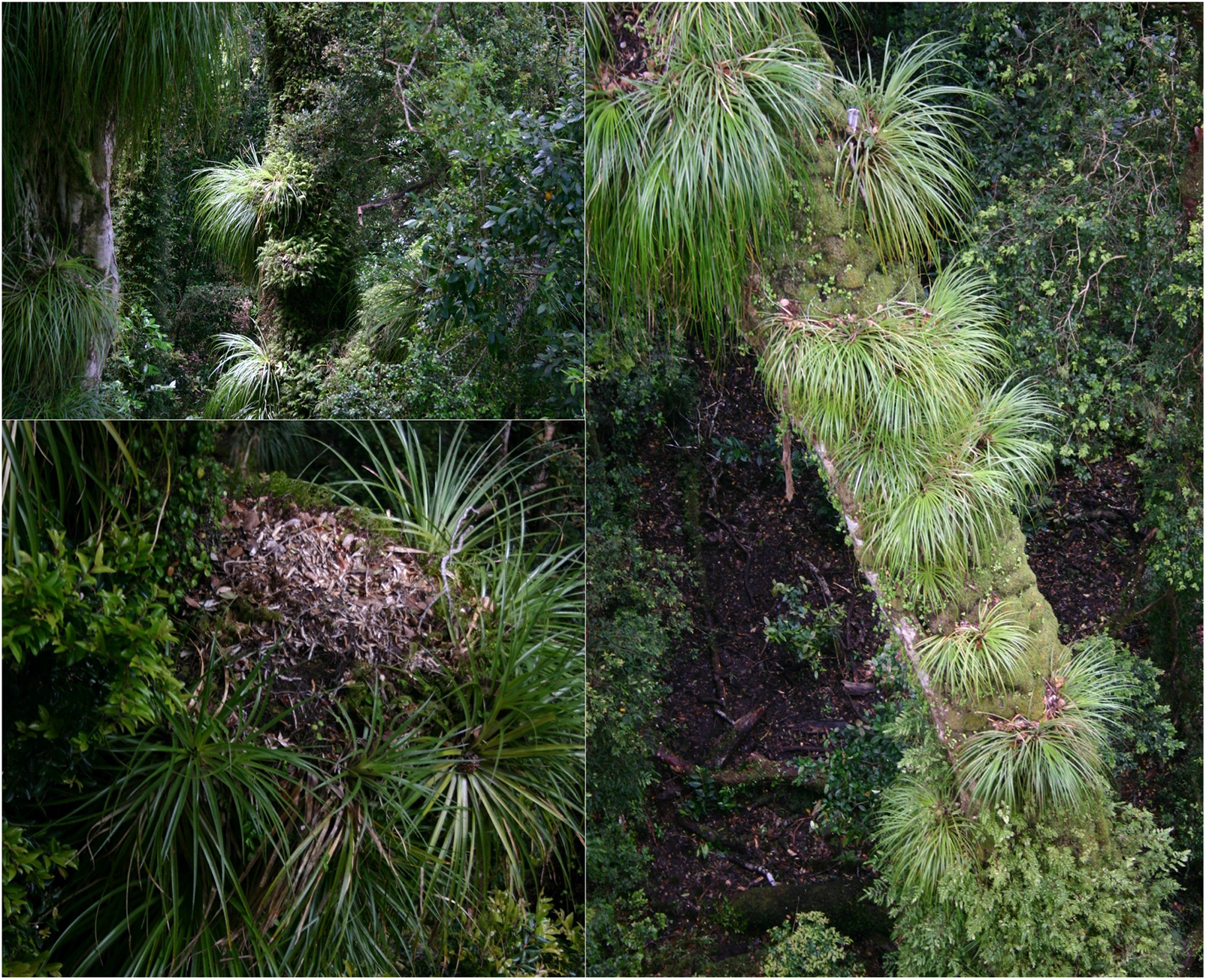
Mean relative interaction index and confidence intervals for soil invertebrate families found in litter and soil samples of *Asplenium nidus* (Beaulieu *et al*. 2010) and *Fascicularia bicolor* (Ortega-Solís *et al*. 2017).

**Figure 5:**
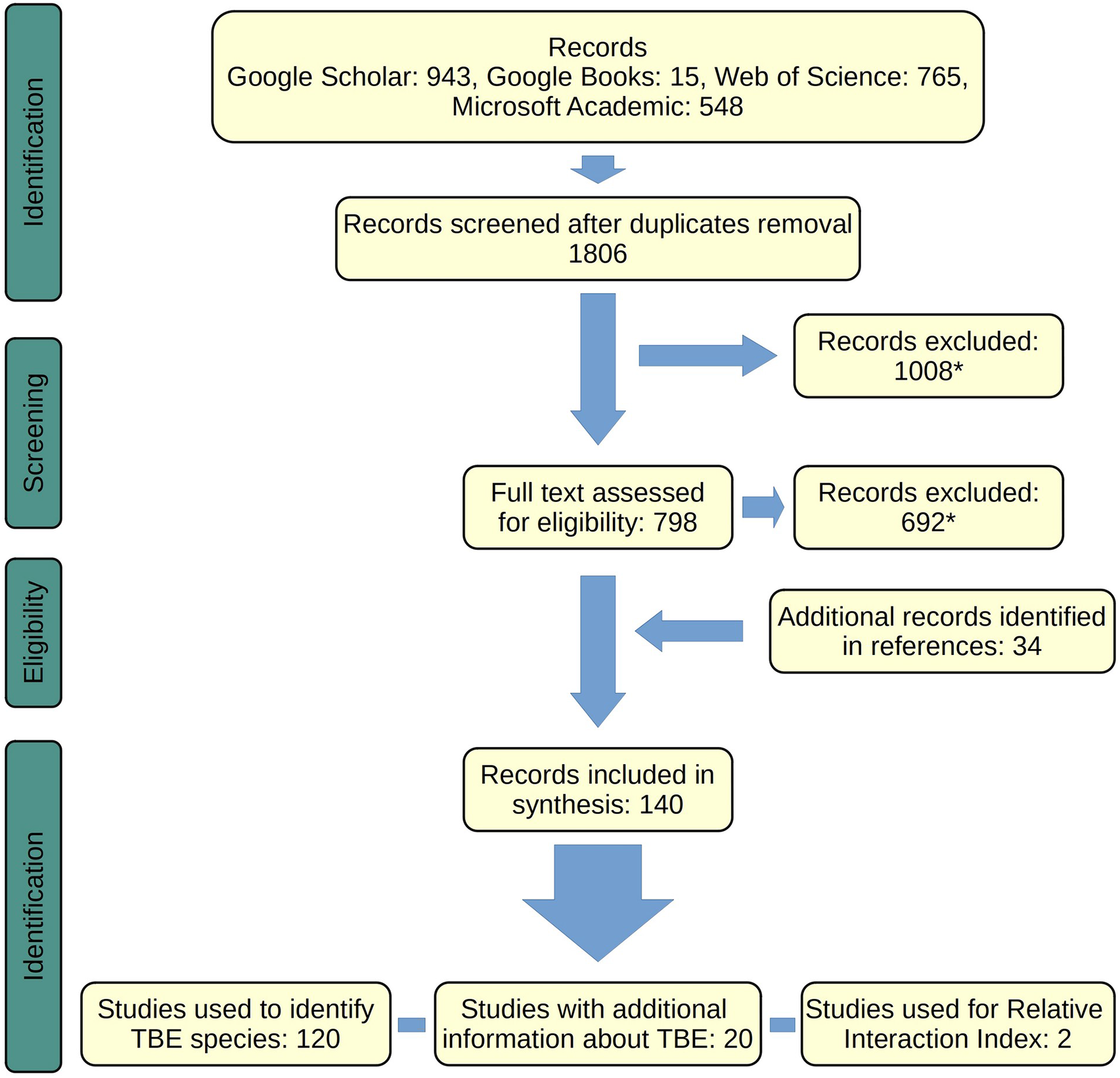
Hypothesized pathways for the impacts of TBE on tree-level biodiversity. **A**: In a tree without epiphytes (or with just a few ones) the colonization of a TBE could create habitats different from those already present on the tree, thus, increasing species richness at the tree level. **B**: If the addition of TBE does not create habitats different from those presented on the host-tree – or if the host already has an important amount of epiphytic material – the abundance of tree dwelling plants and animals could increase because of the greater amount of habitat available without necessarily increasing species richness. **C:** If different TBE species colonize a tree, they could increase the abundance of individuals and the species richness on the tree because they provide additional habitat, with each TBE species creating a slightly different habitat (modified after Angelini and Silliman 2014). Images from <u>Openclipart</u>.

**Table 3:**
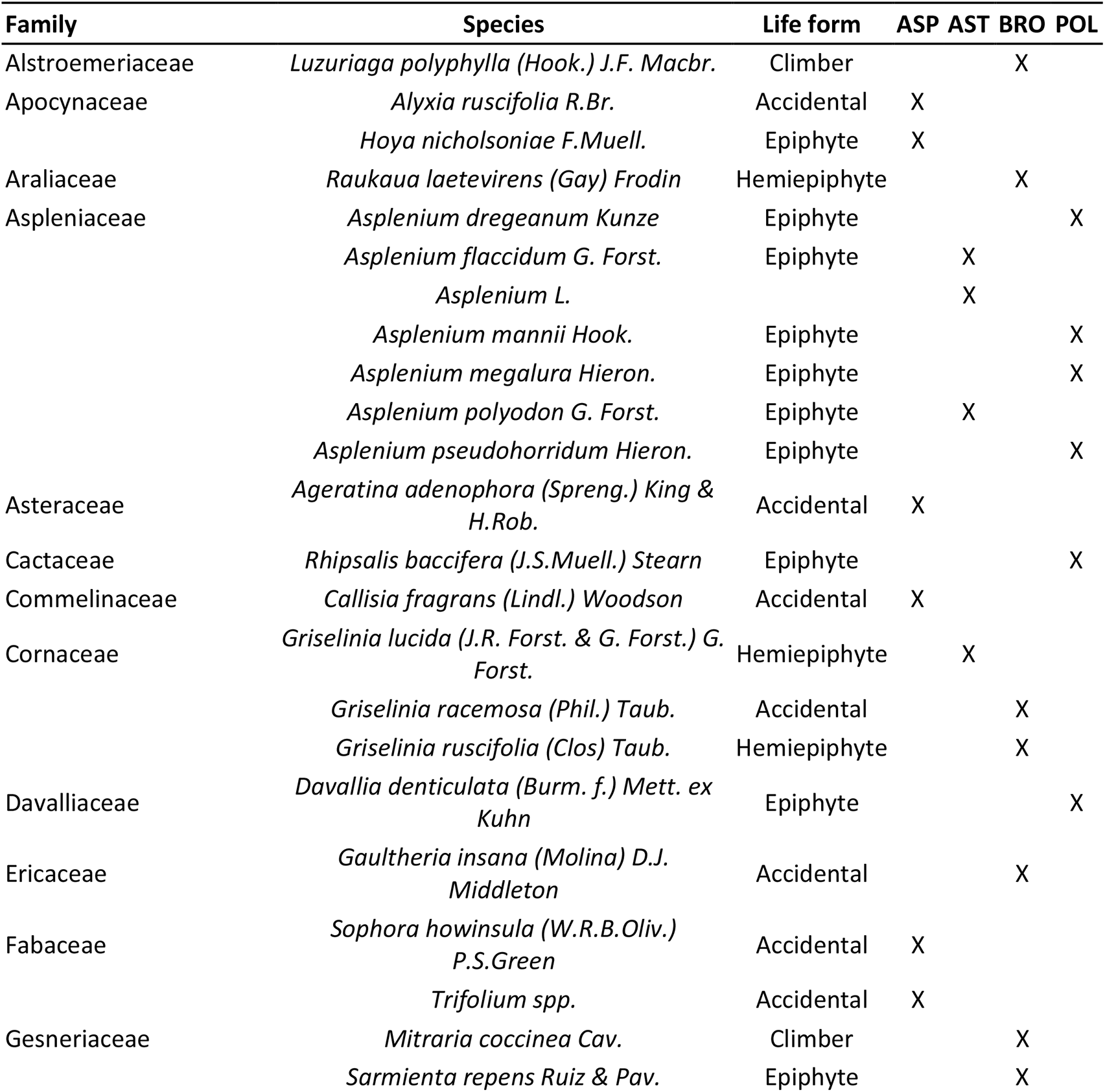
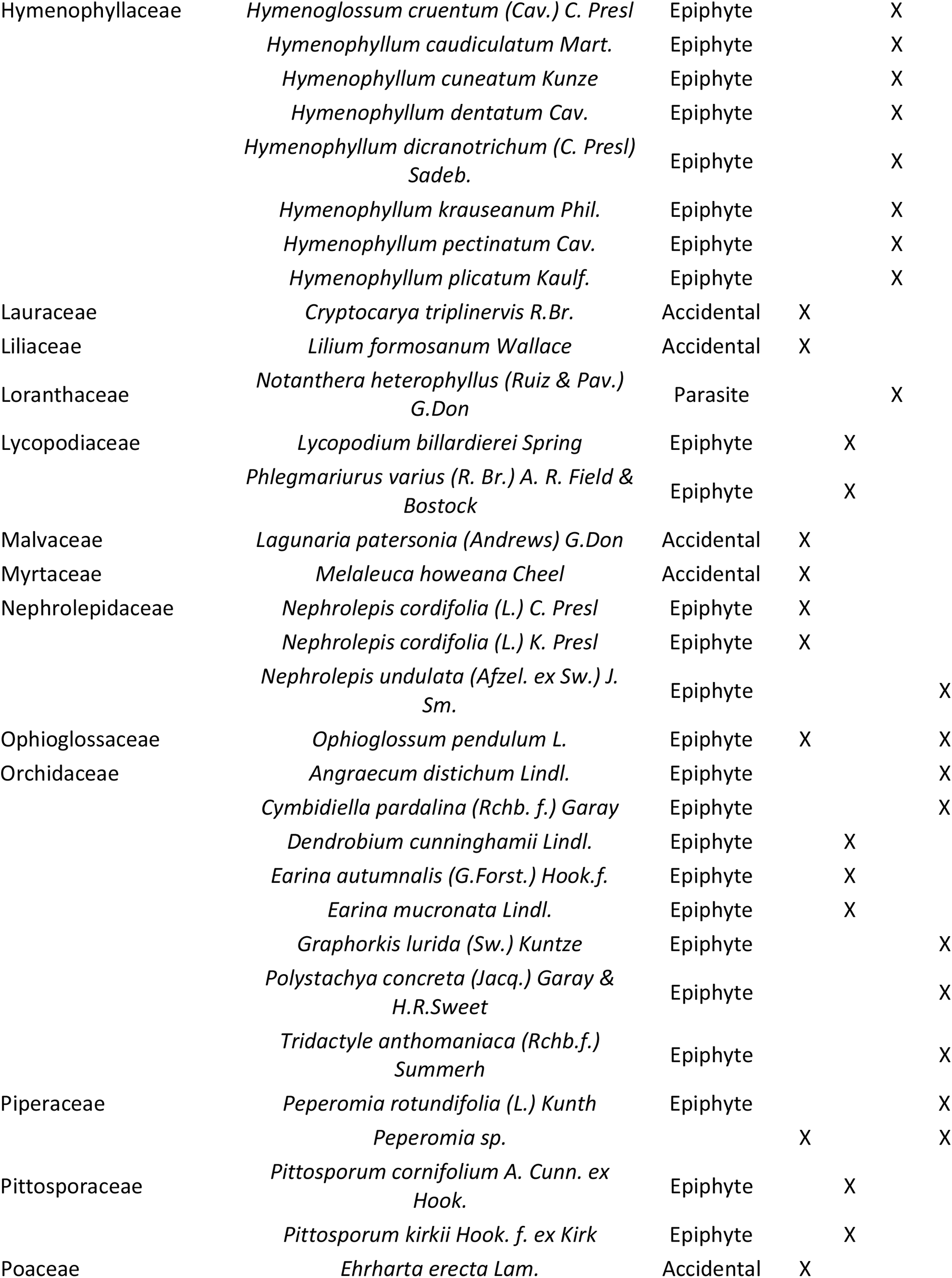
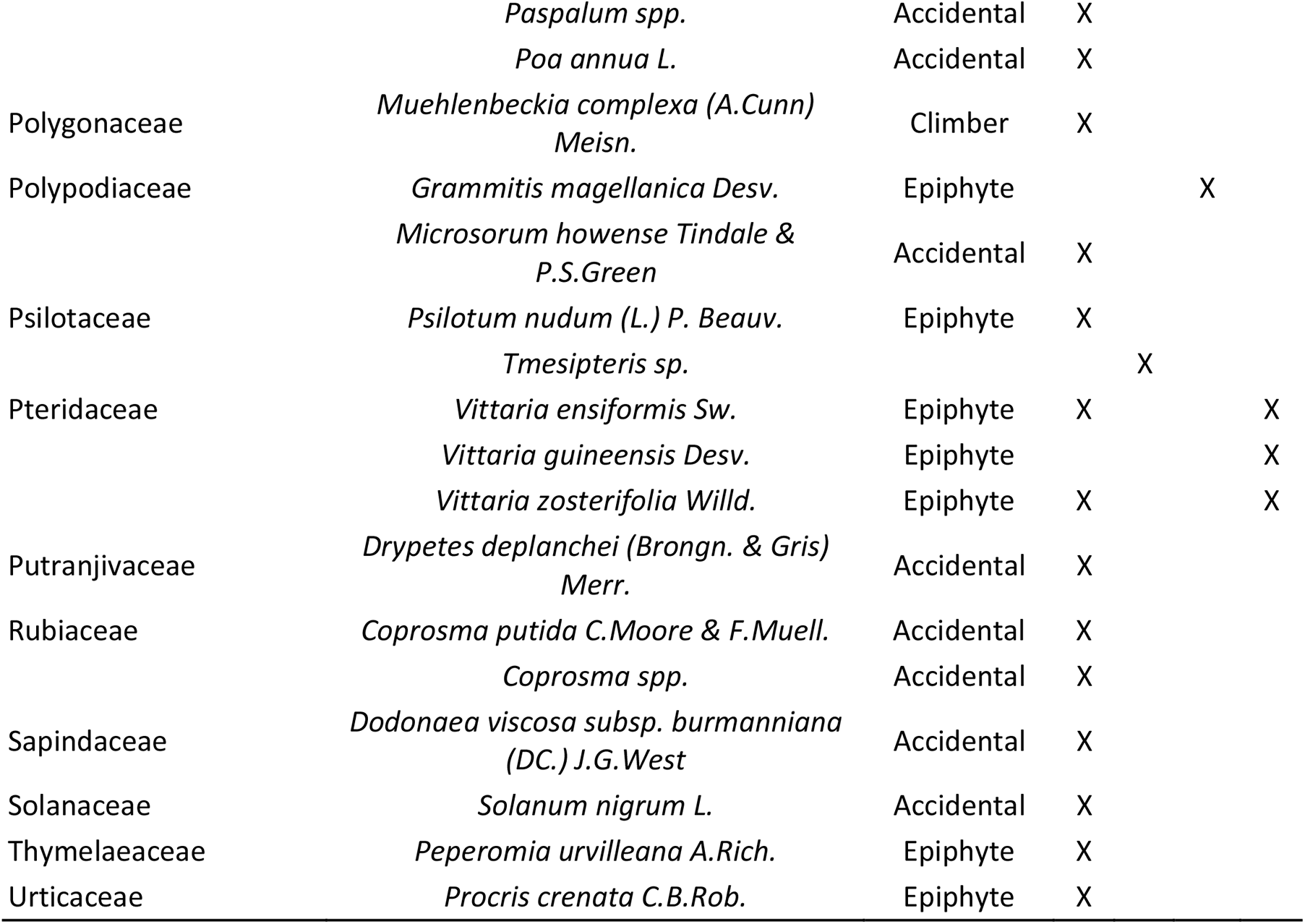
List of plant species that thrive within TBE. *Indicates genus-level identification. Life form categories were taken from Lawalrée (2000), Breitwieser *et al*. (2010) Rodriguez *et al*. (2018), and Zotz *et al*. (2021). TBE families where each species was found were **ASP**: Aspleniaceae, **AST**: Asteliaceae, **BRO**: Bromeliaceae, and POL: Polypodiaceae. A detailed list of references per species is provided as Supplementary Information.

The invertebrate communities of TBE seem to be dominated by generalist species of soil dwellers that can also be found in arboreal and terrestrial soils not associated with TBE (Ortega-Solís *et al*. 2017) and differ from invertebrates inhabiting the leaves and bark of the host trees (Ellwood and Foster 2004). The influence of TBE on invertebrate diversity is likely to be driven by niche construction and cascading trophic effects. The TBE increase the habitat available for soil invertebrates in the canopy by enhancing arboreal soil accumulation in forks and horizontal branches, or even by becoming attached to vertical surfaces that do not accumulate soil (Ortega-Solís *et al*. 2017). Moreover, because most invertebrates are poikilotherms (Meehan 2006), they also benefit from the amelioration of temperature extremes within the TBE microhabitat in the forest canopy. Similar to the communities in tank bromeliads (Richardson 1999; Balke *et al*. 2008; McCracken and Forstner 2014), TBE support a specialized food web including detritivores (i.e., oribatid mites) and predators (i.e., spiders, amphibians, and reptiles), and possibly exert a bottom-up control where the basic resource that sustains invertebrate populations is the litter and other debris trapped by TBE. Detritivores are directly dependent on the litter retained by TBE and contribute to litter fragmentation, enhancing decomposition and nutrient release. In turn, they can be preyed upon by spiders and myriapods that can be also preyed on by larger predators (i.e., spiders, amphibians, reptiles, and birds). The microcosm of TBE seems to be an under-explored model system suitable for testing the influence of species composition of the invertebrate community on ecological functions. For instance, clumps of *Collospermum hastatum* has been used as an island surrogate to model the effects of island size on the invertebrate community and nutrient cycling (Wardle *et al*. 2003). That study shows that larger TBE clumps retain more nitrogen and phosphorus and have a higher density of macro- and micro-arthropods than smaller clumps, while the density of Diplopoda, Isopoda, and fungal-feeding nematodes decrease (Wardle *et al*. 2003). Since invertebrate decomposers are actively involved in nutrient cycling, decreased nutrient retention could be driven by changes in the decomposer community (Wardle *et al*. 2003). Species composition and abundance of oribatid mites inhabiting nest fern TBE are also affected by fern size (Karasawa and Hijii 2006a). Thus, changes in nutrient cycling driven by changes in the composition of the invertebrate community are likely to happen across TBE species.

Invertebrate abundances and biomass are probably the biodiversity parameters most affected by TBE. Invertebrates inhabiting TBE are not unique to them, yet are shared with forks, branches, or non-TBE species where litter and arboreal soils accumulate. Thus, the diversity of invertebrates is less likely to be affected by the presence of TBE if they do not contribute a high percentage of the epiphytic material or if few invertebrate species are unique to TBE. However, the total abundance and biomass of invertebrates – in the host tree and the forest stand– could still increase because of the additional habitat provided by TBE. Nest ferns in lowland forests of Borneo contain ca. 3.77 kg ha^-1^ of invertebrate biomass, while the same estimate for trees only is 3.82 kg ha^-1^ (Ellwood and Foster 2004). Biomass increases exponentially with the size of TBE individuals (Ellwood and Foster 2004), and therefore, larger ones are likely to support more invertebrate biomass than expected from simple linear extrapolation. Moreover, most studies regarding TBE-invertebrate associations are based on a taxonomic approach, while a functional diversity approach (i.e., based on traits) remains unexplored. Therefore, we suggest that future studies should consider functional diversity to assess the underlying mechanisms of diversity changes associated with TBE and the varying amount of microhabitat they provide.

Amphibians, reptiles, birds, and mammals also use TBE, yet the potential importance of TBE as a resource or microhabitat for vertebrates is virtually unknown. Amphibians and reptiles are poikilotherms that benefit from the microclimatic control exerted by TBE (Scheffers *et al*. 2014; Seidl *et al*. 2020). Even large reptiles can benefit from TBE. For instance, the python *Simalia kinghorni* dwells on *Drynaria rigidula*, *Platycerium bifurcatum,* and *Asplenium* TBE 17 to 40 m above the forest floor during the cold season in Australia (Freeman and Freeman 2009). *S. kinghorni* individuals frequently return to the same TBE (Freeman and Freeman 2009) and Australian aborigines have been reported to hunt them in *“large clusters of ferns found on the trunk of trees’’ (Lumholtz and Anderson 1889)*, which suggests that TBE are important habitats for these snakes. Bats from the genus *Cynopterus* sometimes have been reported building roosting sites in *A. nidus* or the orchid *Cymbidium finlaysonianum*. Moreover, a study of roosting habits of the bat *Balionycteris maculatus* found that 25% of the roost sites were in the rootball of *A. nidus* (Hodgkison *et al*. 2003). Multiple bird species use TBE as nesting sites, even returning to them annually (Barea 1997; Roland *et al*. 2005). However, most of the species described also use other nesting substrates and thus are not uniquely dependent on TBE.

Moreover, TBE facilitate the establishment of other epiphytes in the forest canopy, but studies trying to explain the ecological mechanisms of such plant-plant interactions are few. We found 29 species of plants inhabiting Aspleniaceae, 18 that live within Polypodiaceae, 17 within the bromeliad *F. bicolor*, and 12 within Asteliaceae species. However, this information could be limited by a general lack of attention on plant-plant interactions involving TBE. Plant families with more species that live within TBE are Hymenophyllaceae and Orchidaceae (11% each). At the species level, the most frequently reported TBE inhabitants are *Griselinia lucida* (4 records), *Ophioglossum pendulum* (4), *Asplenium polyodon* (3), *Pittosporum cornifolium* (3), and *Vittaria zosterifolia* (3). TBE facilitate other plants by providing a thick arboreal soil layer where they can root, and readily access nutrients and water (Jian *et al*. 2013; Ortega-Solís *et al*. 2017). Field observations of plants inhabiting *F. bicolor* clumps suggest that habitat partitioning occurs within these micro-refugia, with woody plants growing on top and filmy ferns and pendant epiphytes growing below (Ortega-Solís pers. obs.). Epiphytes established in the upper section of TBE could require deeper soils or greater light exposure, while those located below TBE could benefit from a darker and less exposed habitat that reduces evapotranspiration. For instance, *V. zosterifolia* is a pendant fern that grows alone or in the rootball of *A. antiquum*; however, the frond length of ca. 60% of individuals inhabiting *A. antiquum* is between 121 and 160 cm while most of those growing alone reach only 40 to 80 cm length (Jian *et al*. 2013). Nutrient concentration does not differ between solitary *V. zosterifolia* individuals and those associated with *A. antiquum* (Jian *et al*. 2013) and they did not significantly increase their biomass in response to fertilization (Huang and Lin 2016). Therefore, the larger size of *V. zosterifolia* co-occurring with *A. antiquum* seems to be driven by greater water availability and reduced evapotranspiration rather than by nutrient supply. Another plant-plant interaction mediated by water availability could occur between *F. bicolor* and filmy ferns. The latter are hygrophilous species (Dubuisson *et al*. 2009) with one-cell layer laminae and no stomata, which make them highly dependent on atmospheric humidity to avoid desiccation. Filmy ferns can inhabit from the base of the trunks to the upper canopy (Clement *et al*. 2001; Krömer and Kessler 2006; Díaz *et al*. 2010), but individuals inhabiting below *F. bicolor* mats have frequently larger laminae and are better hydrated than those growing alone even during hot and dry summers (Mellado-Mansilla pers. obs.; see Figure 2 in Supplementary Information).

The occurrence of bacteria, fungi, and other microorganisms in TBE and arboreal soils also remains mostly ignored. Gram positive and gram negative bacteria had the highest relative importance in *Asplenium nidus*, *A. phyllitidis*, and *A. serratum*, ranging from 19 to 21%, followed by fungi (5-11%) and actinomycetes (4%) inferred from the analysis of phospholipid fatty acid profiles (Donald *et al*. 2017, 2020). A comparison between *A. phyllitidis* and *A. serratum* (from Borneo and Amazonia respectively) show they have the same groups of microorganisms but different compositions, with a greater abundance of fungi in the drier conditions of Borneo (Donald *et al*. 2017, 2020). Fungi are more likely to succeed in the canopy of dry forests because they are able to transfer water through their hyphae (Guhr *et al*. 2016; Donald *et al*. 2017). Fungi also form litter-trapping networks on their own (Snaddon *et al*. 2012). Despite not having found studies about litter-trapping fungi inhabiting TBE, such an association may be a common occurrence. Aquatic fungi also inhabit *Drynaria quercifolia* in riparian zones of south west India (Sridhar *et al*. 2006; Karamchand and Sridhar 2009). Reports of protozoa and other microorganisms inhabiting TBE are lacking. Since microorganisms are fundamental in nutrient cycling, we suggest that studies about the diversity of micro-organisms in arboreal soils should be encouraged.

At the tree scale, TBE may affect biodiversity in a number of ways (Fig. 5). Trees without a significant accumulation of epiphytic species benefit from the establishment of a few TBE individuals, which increase microbial and invertebrate species diversity by creating new habitats that differ from those offered by the tree alone (Figure 5A). If the host tree already has arboreal soil or epiphytes, its colonization by TBE individuals should increase the abundance of invertebrates and plants requiring arboreal soil, but may not increase species richness (Figure 5B). If different TBE species colonize the same tree, they should increase the abundance and species richness of soil invertebrates and soil-rooted plants, assuming that every TBE species will create slightly different habitats (Figure 5C). It should be noted that beyond the direct effects proposed in Figure 5, other indirect effects could happen. For instance, *Crematogaster difformis* ants nesting in *Platycerium* and *Lecanopteris* ferns (the latter is a mirmecophytic fern, but is not a TBE) have been shown to remove lianas from their territory (Tanaka and Itioka 2011). On the other hand, some ant species build gardens by bringing seeds to their nests (Morales-Linares *et al*. 2018) or providing shelter to mirmecophylous invertebrates (Maruyama 2010; Maruyama *et al*. 2014). In summary, the influence of ants on the biodiversity found on trees exemplify the cascading effects that could emerge from TBE and their associated species.

## RELEVANCE OF TRASH-BASKET EPIPHYTES FOR FOREST MANAGEMENT AND CONSERVATION

The rapid loss of forest biodiversity is a major concern for conservation biology. Forest degradation has continued to increase over the past decades, despite efforts to reduce it (Hansen *et al*. 2013). Even if forest regrowth is expected in some landscapes, the recovery process of canopy biodiversity is an unresolved challenge. In this context, restoring TBE could be useful for enhancing or restoring canopy species diversity and their associated ecosystem functions. Three key caveats must be addressed in this regard:

**1) In which ecosystems can TBE have the greatest influence?** The relative importance of TBE in different forest ecosystems remains to be explored. As well as cushion plants, the influence of TBE on biodiversity and ecosystem functions depends on the amount and uniqueness of the habitat they create. If epiphytic litter and arboreal soil are abundant in the forest canopy, increasing TBE populations likely would not have detectable effects, because the accumulation of arboreal soil is driven by other factors. On the other hand, TBE could have significant impacts in secondary forests lacking epiphytic plants or in tree plantations, where biotic homogenization, distance to propagule sources, or short harvest cycles would otherwise prevent the formation of a fully-fledged community within the canopy.
**2) Which species benefit from TBE?** Whether conservation or the active establishment of TBE in forests will produce positive effects on forest biodiversity depends on the species we expect to conserve, or the ecosystem functions we would like to restore. *A. nidus* individuals occurring in oil palm plantations provide microhabitat conditions similar to those in natural forests and support almost the same levels of ant species richness (Fayle *et al*. 2010). However, the composition of the ant community supported by *A. nidus* in plantations is dominated by exotic species (Fayle *et al*. 2010). Exotic species can have detrimental effects on native species (i.e., through competition), but they could also have positive outcomes for the ecosystem if they provide similar functions as locally extinct or rare native species.
**3) Is there sufficient knowledge about the reproduction and establishment of TBE to develop viable restoration plans?** TBE such as *Asplenium* and *Platycerium* ferns are cultivated and sold in garden stores, which means there could be enough practical knowledge to propagate them for restoration efforts. Despite being less common, *Fascicularia* and *Anthurium* have also been cultivated by gardeners or maintained in greenhouses, suggesting that the knowledge gap to reproduce them at larger scales may be smaller than for other TBE. However, maintaining the genetic diversity of cultivated plants or restored assemblages could be challenging. *Fascicularia* reproduces clonally and new individuals are commonly grown from rosettes rather than from seeds. *Anthuriums* can be reproduced from seeds or from cuttings, the latter producing clones instead of genetically distinct individuals. Also, *F. bicolor* likely disperses via endozoochory, but the identity of its dispersers is unknown. If the seed dispersers do not occur in a restoration site, the long-term persistence of *F. bicolor* as epiphyte in the restored area may be uncertain.

Many TBE are often associated with large, senescent trees in old-growth forests (Cummings *et al*. 2006; Fayle *et al*. 2009; Ortega-Solís *et al*. 2020) and thus they are less likely to to establish in young secondary forests. *A. antiquum* grows in forest stands with closed canopies, while *A. nidus* colonizes tall canopy trees and occupies sun-exposed locations (Fayle *et al*. 2009). *A. nidus* also predominates in oil palm plantations (Turner and Foster 2006), thus it seems more resilient than *A. antiquum* to environmental changes. To advance in the inclusion of TBE in restoration efforts, more studies of their functional ecology are required to assess their responses to water stress, temperature, light, and nutrient levels in the canopy, as well as additional studies to document the abundance of individuals and their host trees in forest ecosystems.

## CONCLUDING REMARKS

The concept of TBE unites a group of secondary foundation species that influence biodiversity and ecosystem functions related to organic matter distribution and accumulation in the forest canopy. Therefore, TBE have considerable potential to contribute to the maintenance of species diversity and ecological processes on the treetops of native forests impacted by human activities or even within managed tree plantations. However, the inclusion of TBE in plans to conserve or restore forest biodiversity is hampered by the lack of knowledge of the ecology, diversity, and habitat preferences of TBE. The specific biotic and abiotic interactions of each TBE species could be also interesting from a theoretical viewpoint, as models to evaluate hypotheses on niche theory, island biogeography, and the links between biodiversity and ecosystem functioning. We enumerated the considerable opportunities and need for further studies of basic ecology and conservation of TBE, which we hope will be addressed in the coming years.

## ACKNOWLEDGEMENTS

We are grateful to M. Sundue, H. Brent, M. Kessler, and Ferns of the World (www.fernsoftheworld.com) for pictures used in Figure 2 and Supplementary Information.

## DATA AVAILABILITY

The data and scripts underlying this article are available in Göttingen Research Online repository, at https://doi.org/10.25625/TANQL6

---

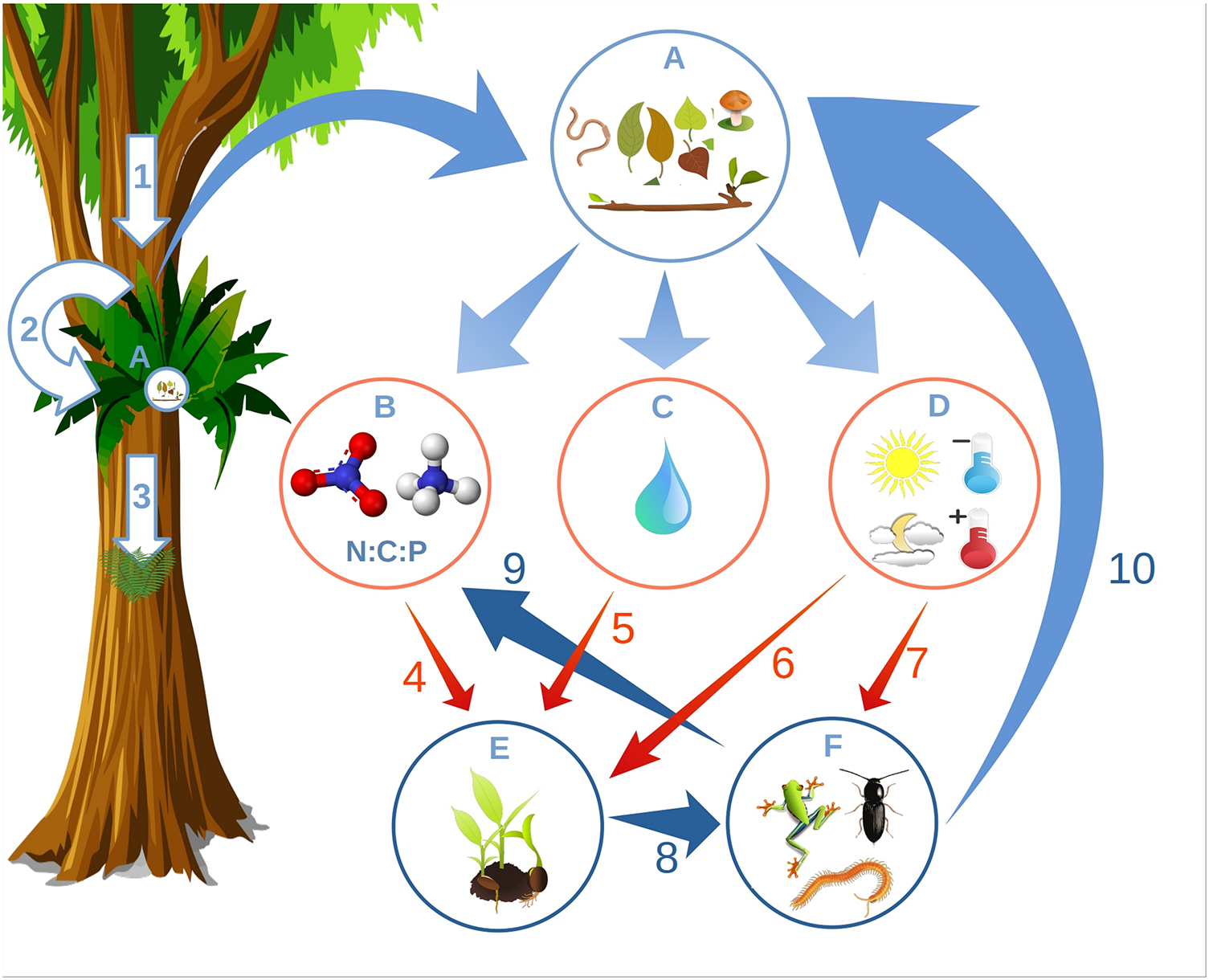

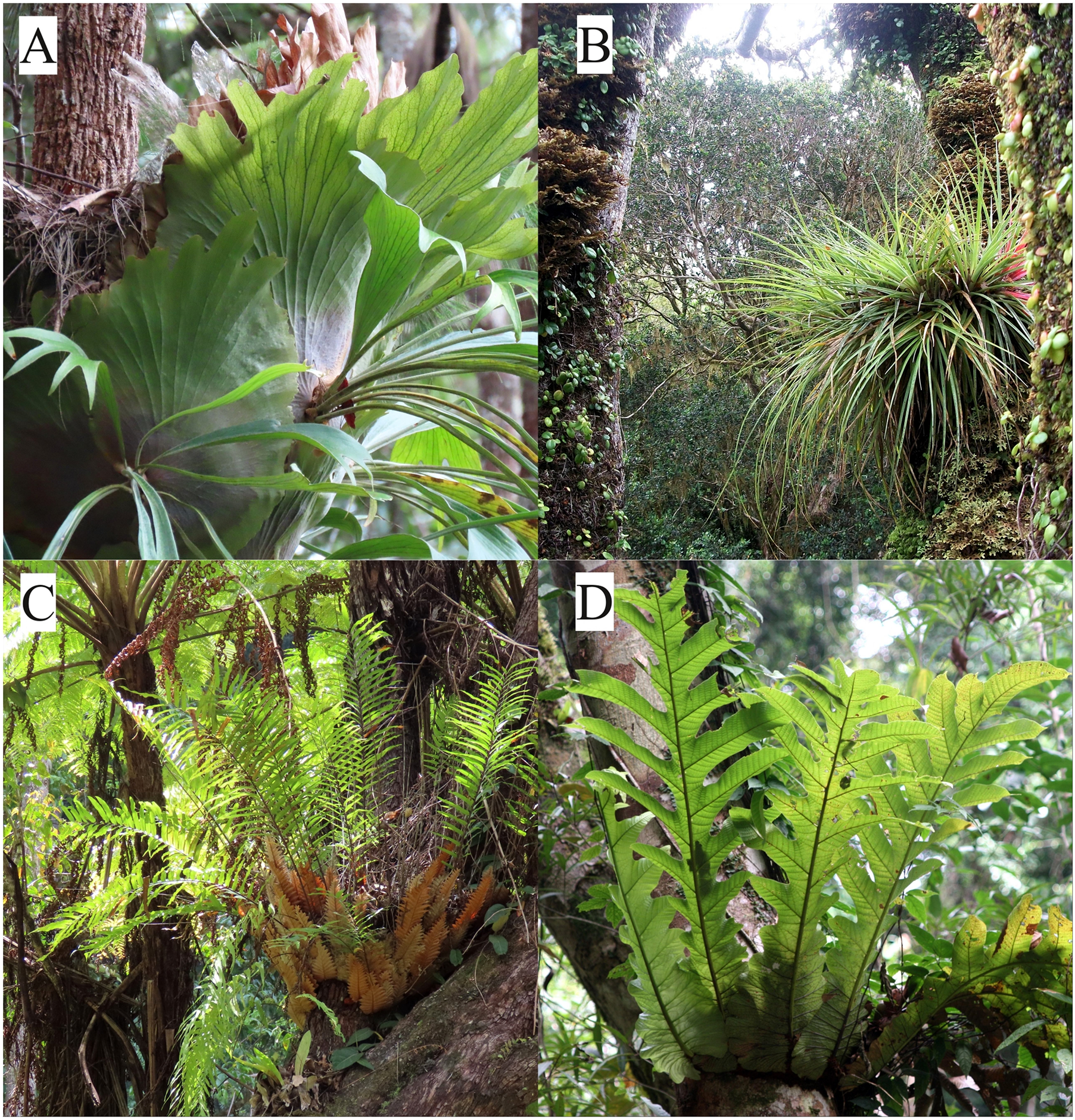

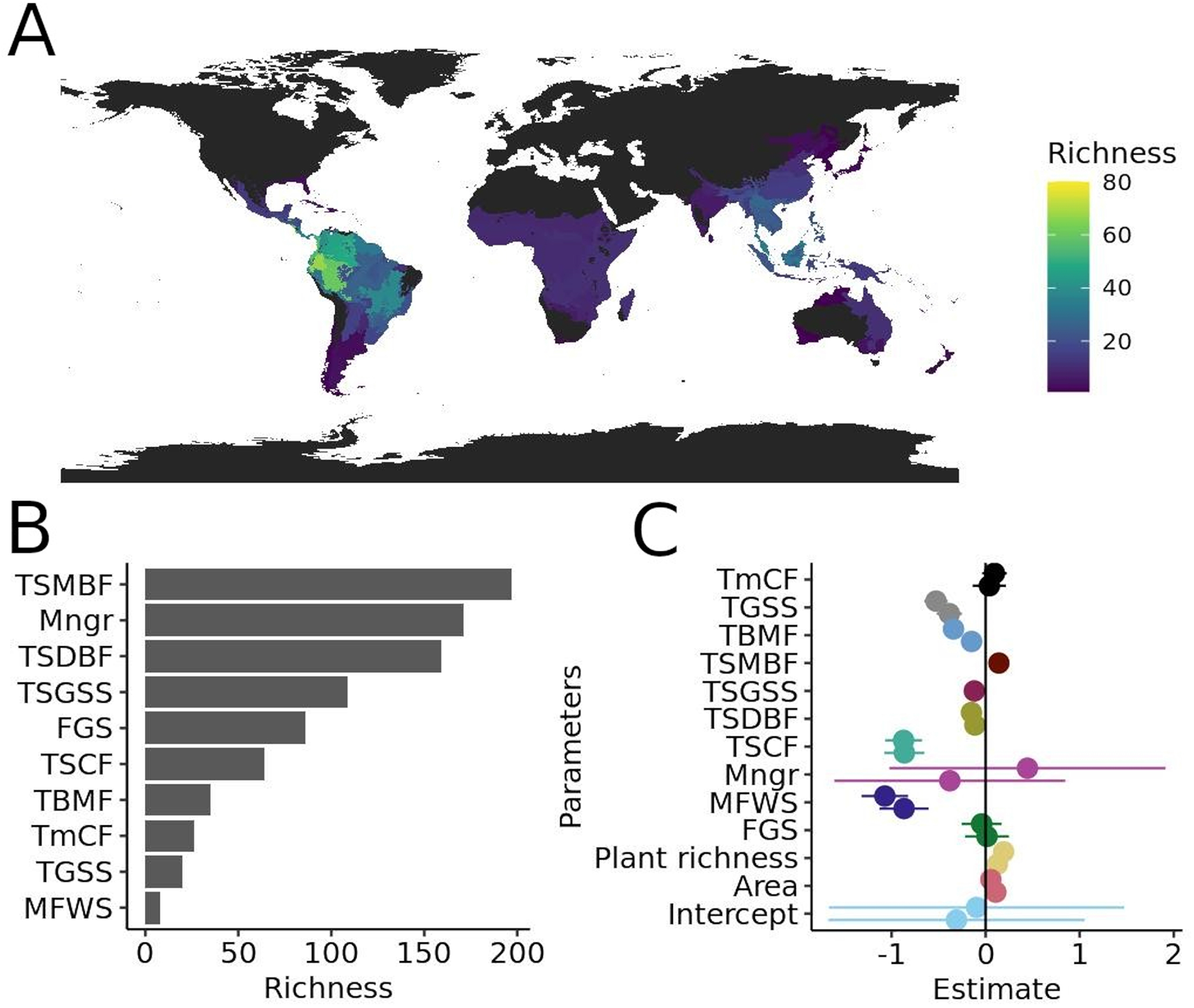

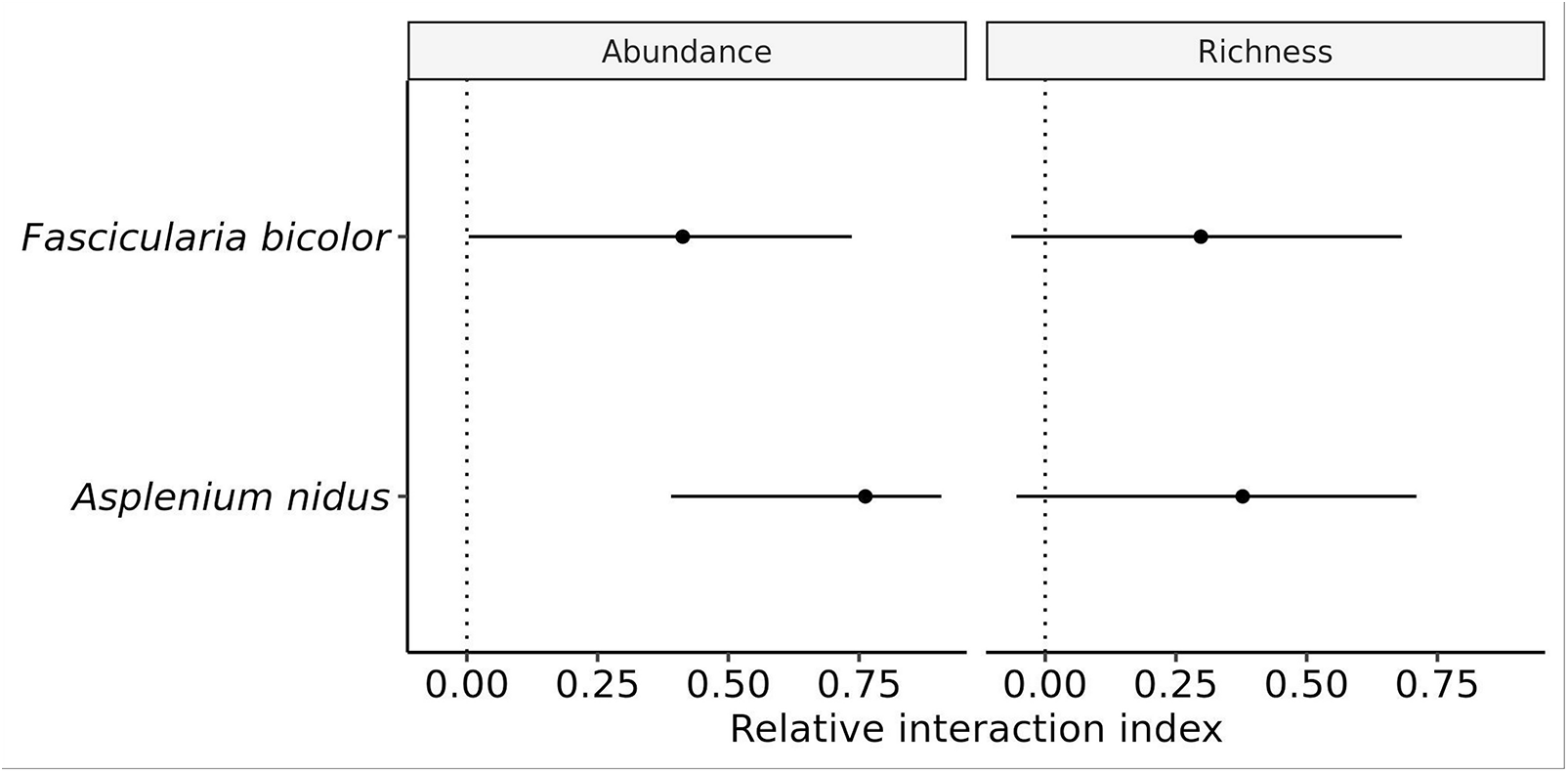

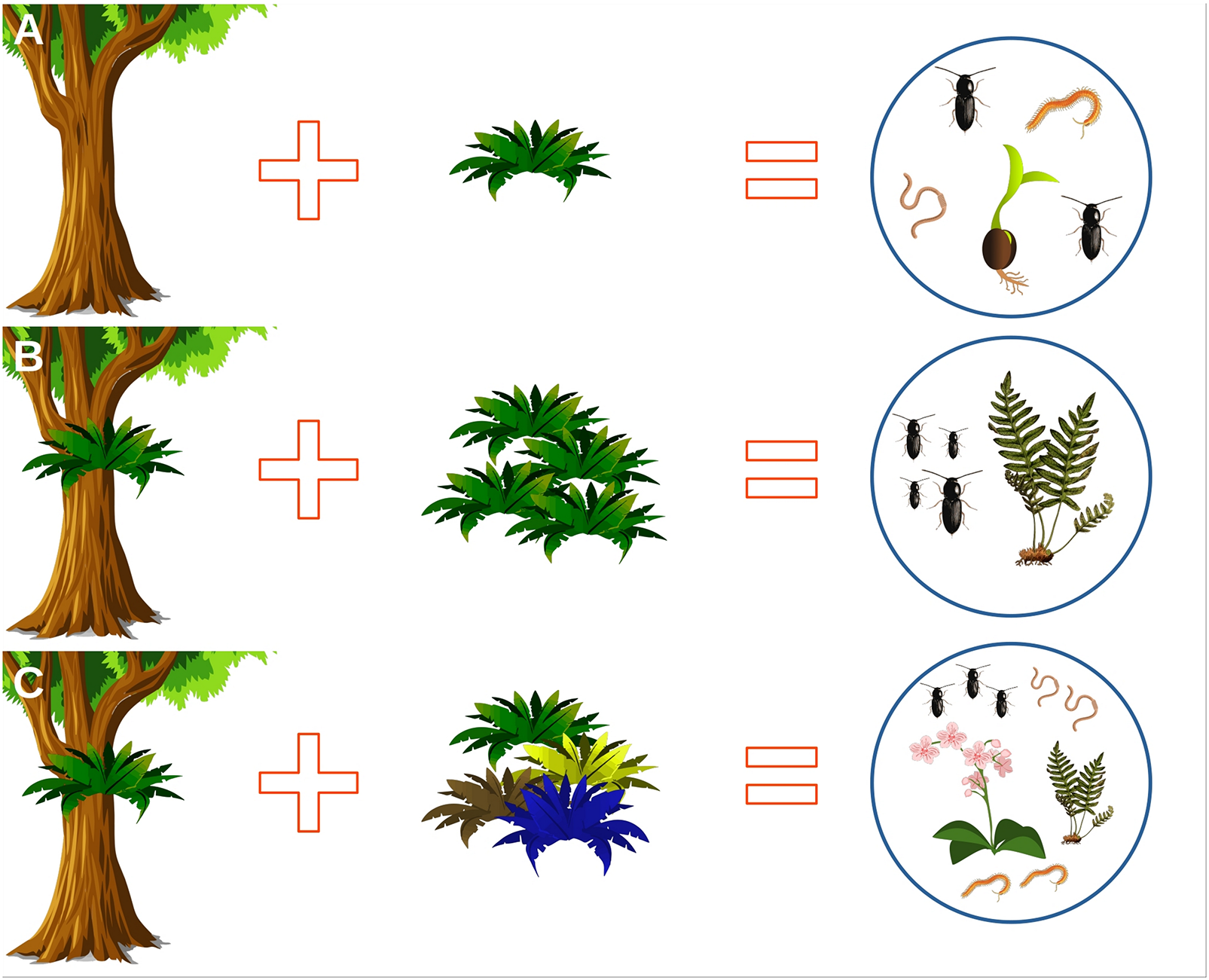

